# Transcription factor enrichment analysis (TFEA): Quantifying the activity of hundreds of transcription factors from a single experiment

**DOI:** 10.1101/2020.01.25.919738

**Authors:** Jonathan D. Rubin, Jacob T. Stanley, Rutendo F. Sigauke, Cecilia B. Levandowski, Zachary L. Maas, Jessica Westfall, Dylan J. Taatjes, Robin D. Dowell

## Abstract

Detecting differential activation of transcription factors (TFs) in response to perturbation provides insight into cellular processes. Transcription Factor Enrichment Analysis (TFEA) is a robust and reliable computational method that detects differential activity of hundreds of TFs given any set of perturbation data. TFEA draws inspiration from GSEA and detects positional motif enrichment within a list of ranked regions of interest (ROIs). As ROIs are typically inferred from the data, we also introduce *muMerge*, a statistically principled method of generating a consensus list of ROIs from multiple replicates and conditions. TFEA is broadly applicable to data that informs on transcriptional regulation including nascent (eg. PRO-Seq), CAGE, ChIP-Seq, and accessibility (e.g. ATAC-Seq). TFEA not only identifies the key regulators responding to a perturbation, but also temporally unravels regulatory networks with time series data. Consequently, TFEA serves as a hypothesis-generating tool that provides an easy, rigorous, and cost-effective means to broadly assess TF activity yielding new biological insights.

## 2 Introduction

Transcription factors (TFs) are DNA-binding proteins that regulate transcription. When a cell is challenged by a change in the environment, it responds by altering the activity of one or more TFs. TFs, through transcriptional changes, are then responsible for altering cellular function and ultimately deciding cell fate. Because of their importance in global cellular programs, measuring differential TF activity between two conditions is a readout of high-level cellular biology and provides critical insight when details of the involved cellular processes are not known.

Experimental methods for measuring TF activity have largely focused on measuring protein-DNA binding, typically by chromatin immunoprecipitation (ChIP), resulting in high quality sequence recognition motifs for many TFs[19, 38]. Yet, not all binding sites lead to altered transcription activity[61, 55, 18]. Consequently, many of the approaches to inferring regulation by TFs combine ChIP data or motif hits with measures of gene expression[10, 28]. Relying on gene expression data, however, limits the effectiveness of these approaches. Gene expression assays, such as RNA-seq, are only indirect measures on actual transcription. RNA-seq is a steady state measure of RNA and reflects a combination of transcription and degradation[26, 57, 22]. Furthermore, the steady state nature of RNA-seq limits the response dynamics of the assay[25, 37, 42, 1, 36], as both newly created and long lived RNAs contribute to RNA measurements[49, 47]. Therefore, directly assaying transcription initiation improves on both the positional and temporal resolution when quantifying the activity of regulatory sites.

A large number of high throughput assays either directly or indirectly assay transcription initiation. Nascent transcription assays[16, 35] directly measure *bona fide* transcription, prior to RNA processing. Cap associated approaches, such as CAGE and GRO-CAP, target the 5′ cap of transcripts[4, 15, 58]. Transcription arises from a subset of nucleosome free regions, therefore chromatin accessibility data indirectly informs on the locations of transcription initiation. Likewise, some histone marks have been associated with actively transcribed regions, such as H3K27ac and H3K4me1/3 [11]. In principle, differential signals from these assays inform on the underlying mechanistic activity of TFs[6].

With differential regulatory data, the objective is to infer which transcription factors are causally responsible for the observed changes. With high quality motifs now residing in numerous databases[34, 43, 38], these catalogs can be leveraged to resolve the concurrent activity of many TFs. Historically, detecting motif enrichment in this way relied on sequences being classified into either signal or background and then calculating motif enrichment in signal sequences relative to background[11, 14]. More sophisticated approaches can take advantage of two additional factors: 1) positional information — where the motif is located relative to a region of interest[8, 6] and 2) differential information — the amount of change occurring within that region of interest[45, 12]. Relatively few techniques encode both types of information[39, 52, 24] and these currently provide no easily accessible software package or web-based application.

Our method, which we refer to as transcription factor enrichment analysis (TFEA) draws inspiration from the popular gene set enrichment analysis (GSEA) algorithm[56]. TFEA improves on our previous position based approach[6] and shows performance comparable to the state of the art motif enrichment approaches. Additionally, TFEA can be applied to a number of regulatory data types including PRO-seq, CAGE, DNAse-seq and ChIP-seq. Finally, TFEA is fast, computationally inexpensive, and designed with the user in mind, as we provide an easy to use web interface (https://www.tfea.colorado.edu), a command-line interface, and an importable Python 3 package. TFEA has the potential to become a transformative tool by providing easy downstream analysis aimed at distinguishing temporal and mechanistic details of complex regulatory networks.

## 3 Results

### 3.1 Overview

Conceptually, when a TF is active, it binds to a set of positions within the genome and alters transcription nearby, both at promoters and enhancers. Importantly, this process can both give rise to new transcripts and alter the levels of existing transcripts. Nascent transcription assays show that when a TF is activated, transcripts arise immediately proximal to the corresponding TF motif[1, 6]. In this work we introduce TFEA, which quantifies positional enrichment of TF motifs across an ordered list of regions (Figure 1). The key input into TFEA is a ranked list of regions of interest (ROIs) that typically are obtained independently from each replicate dataset but can also be a list of annotated regions, such as known promoters.

**Figure 1:**
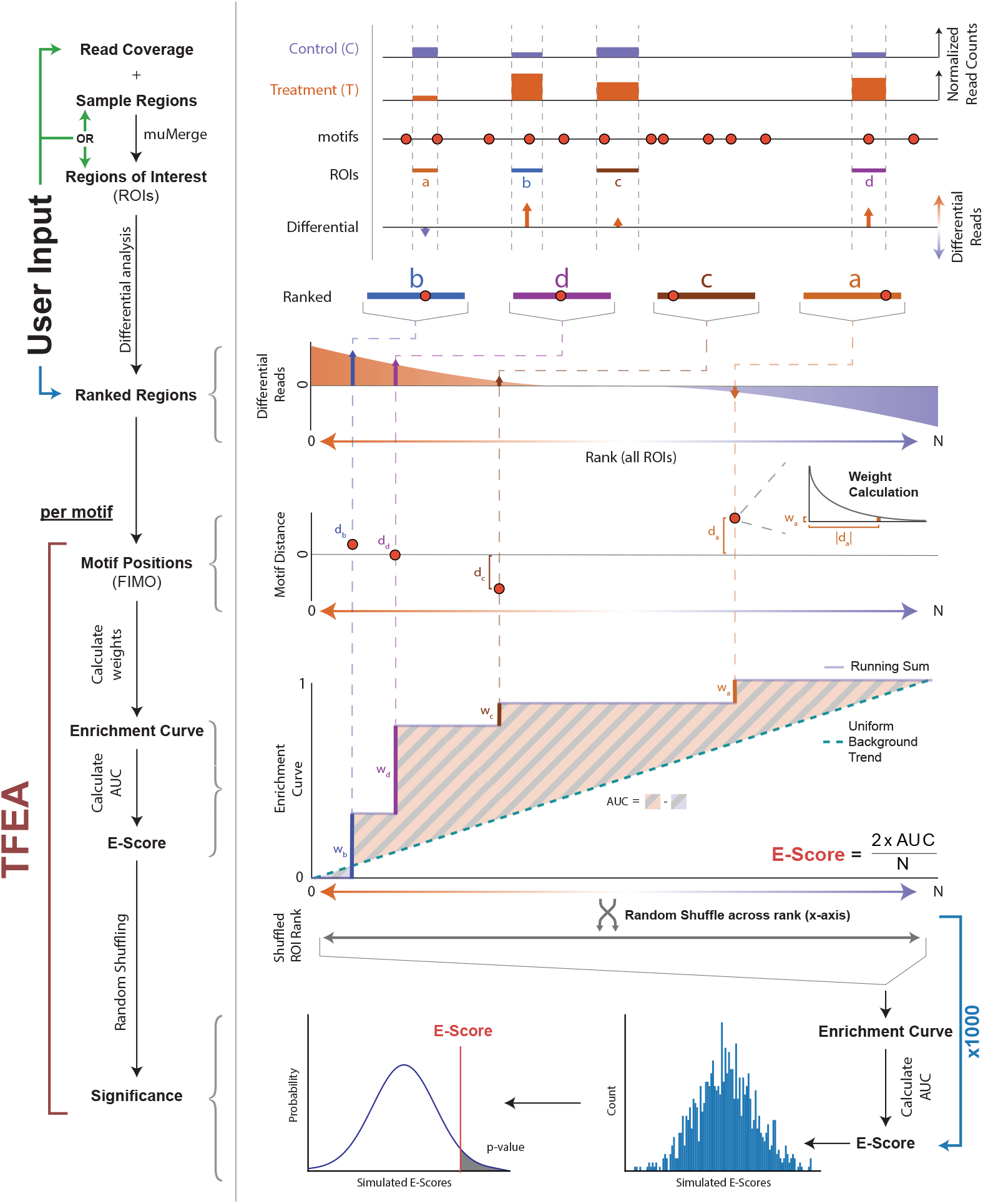
TFEA calculates motif enrichment using differential and positional information. The TFEA pipeline requires, minimally, a ranked list of ROIs. Optionally, a user may provide raw read coverage and regions, in which case TFEA will perform ranking using DE-Seq [2, 41] analysis. With a set of ranked ROIs, TFEA analyzes motif enrichment for each motif provided as a. meme database. For each motif, positions are determined by FIMO scans and an enrichment curve is calculated by weighting each motif instance (using an exponential decay function) and adding this value to a running sum. An E-Score is calculated as 2 * the AUC, e.g. the area under the curve between the running sum and a uniform background (line). For statistical significance, the ROI rank is shuffled 1000 times, and E-scores are recalculated for each shuffle. The true E-Score is then compared to the distribution of E-Scores obtained from the shuffling events. For example output of TFEA see Supp Fig 1 and Supp Fig 2.

### 3.2 *muMerge*: Combining genomic features from multiple samples into consensus regions of interest

A key challenge in defining a set of consensus ROIs is retaining positional precision when combining region estimates that originate from different samples (replicates and conditions). To this end, we developed a statistically principled method of performing this combination called *muMerge* (See Supp. Fig. 3 and online method section 4.1.1 for details). In order to demonstrate the efficacy of *muMerge*, we compare its performance to two common methods for combining regions across multiple samples—merging all samples (e.g. with *bedtools merge*) and intersecting all samples (e.g. with *bedtools interesect*). We performed two tests using simulated data (Supp. Fig.4). For each replicate, we performed 10,000 simulations of sample regions for a single loci, and calculated the average performance.

Using the simulated regions, we first evaluate the methods’ precision as the number of replicates increases. In Fig. 2a, we observe that as the number of replicates increases *muMerge* converges on the correct theoretical loci position (*μ*) more quickly than the other two methods, while still maintaining the correct width for the region (Fig. 2b).

**Figure 2:**
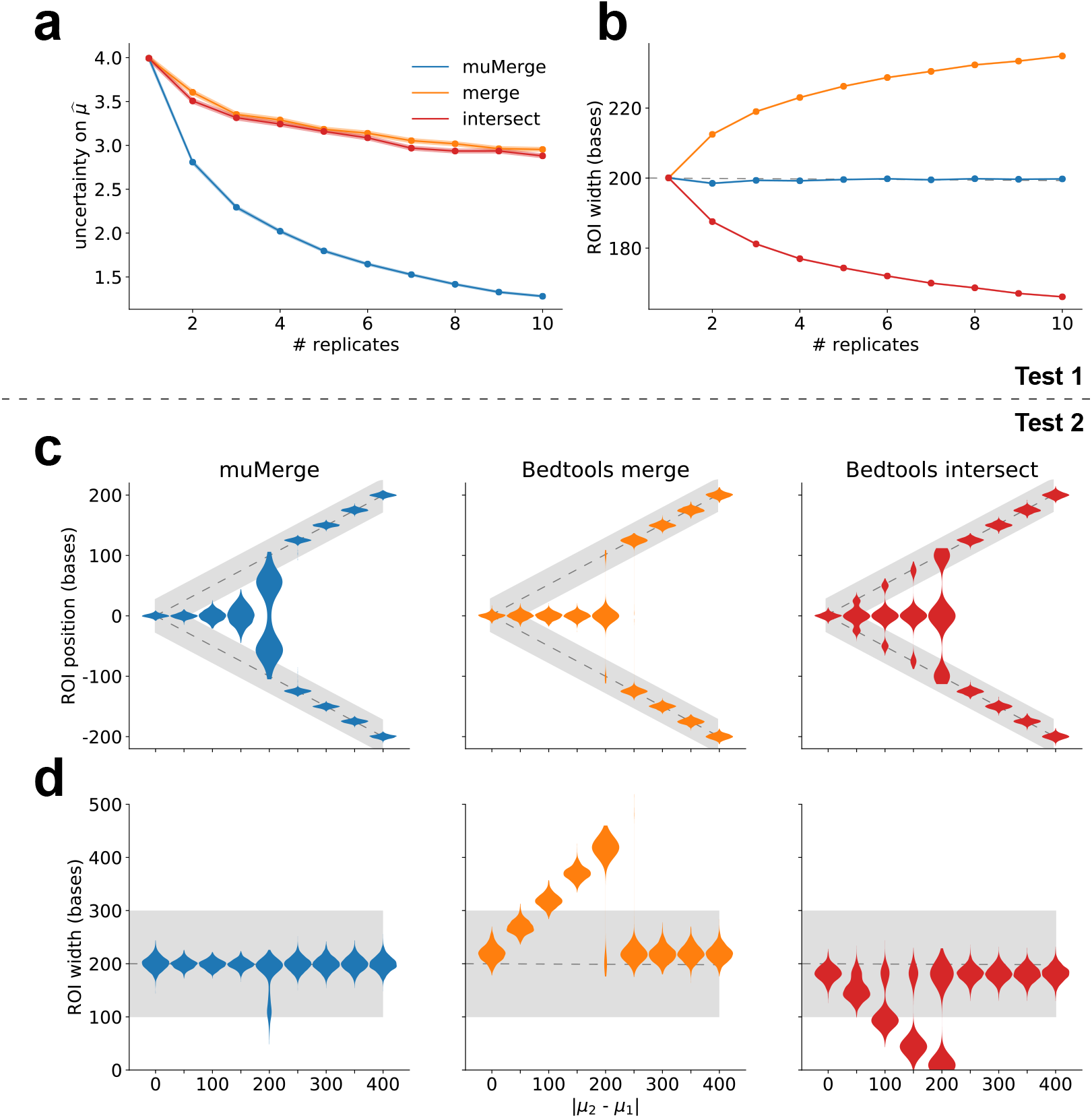
*muMerge* precisely combines multiple samples into consensus ROIs. Here we show a comparison of three methods (*bedtools merge*, *bedtools intersect*, and *muMerge*) for generating ROIs from multiple samples (See Supp Fig 4 for schematic of each method). Test 1 demonstrates the position and width accuracy of a calculated ROI for a single loci, *μ*, as the number of sample replicates are increased (from one to ten). With *muMerge* (a) the positional uncertainty decreases quickly while the (b) estimated ROI width remains relatively constant. Standard error, indicated by shading, is less than the line width. Test 2 demonstrates the precision of the calculated ROI for two closely spaced loci, *μ*_1_ and *μ*_2_, as the spacing between them is increased. In this case, *muMerge* (c) transitions from a single loci to two distinct loci more gradually and (d) the estimated ROI widths do not deviate from the expected value. In all cases, expected value and variance for simulations is indicated by grey lines and shading, respectively. For further detail on the results of Test 1 and 2 and how the simulations were performed, see Supp Fig 4 and methods section 4.3.1.

The second test we performed sought to evaluate the accuracy of these methods when inferring two closely spaced loci, with increasing distance between those loci (Fig. 2c). While closely spaced loci are challenging to distinguish, we observe that *muMerge* smoothly transitions from calling a single inferred loci (when *μ*_1_ and *μ*_2_ are too close to be resolved) to two distinct loci. In contrast, the merge and intersect methods show abrupt transitions that follow increasingly poor ROI width estimates (Fig. 2d).

### 3.3 Transcription Factor Enrichment Analysis

Armed with the defined set of ROIs, the goal of TFEA is to determine if a given TF motif shows positional enrichment preferentially at regions with higher differential signal. Positional enrichment is consistent with the TF contributing to observed alterations. In prior work, we assessed the enrichment of motifs relative to positions of RNA polymerase initiation using a co-occurrence metric referred to as a motif displacement score (MD-Score; see Supp. Fig. 5 for full details)[6]. Unfortunately, the MD-Score approach not only ignored alterations in transcript levels (See Supp. Fig. 6) but also utilized an arbitrary distance threshold that classified motif proximity in a binary fashion.

To include transcript levels into the metric, we can rank ROIs by differential signal (e.g. transcription) before subsequently performing motif displacement calculations within these regions. The simplest approach to this problem is to compare the MD-Scores between the set of differentially transcribed regions and regions whose transcription is unchanged, a method we refer to as the differential motif displacement analysis (MDD, see Supp. Fig. 7 for full details)[52, 24]. Unfortunately, the MDD method introduces an additional arbitrary threshold to classify regions as differentially transcribed or not and still uses the distance threshold set by the MD-Score approach.

In TFEA, we sought a non-binary enrichment metric that accounts for not only the underlying changes in transcription but also the positional enrichment of the motif (Fig. 1). We begin by leveraging the statistically robust, gold standard DE-Seq package[2, 41] to rank regions based not only on the differential p-value but also the direction of fold change. Each region of interest then contributes positively to the enrichment curve in a weighted fashion. These weights are determined by the distance of the motif to the reference point using an exponential function to favor closer motifs. The subsequent enrichment score (E-Score in Fig. 1) is proportional to the integrated difference between the observed and background enrichment curves, calculated as the area under the curve (AUC) in Fig. 1 (see Eq. 8 for precise definition). The background (null) enrichment curve assumes uniform enrichment across all ROIs, regardless of differential signal.

By default, TFEA accounts for the known GC bias of enhancers and promoters by incorporating a correction to the enrichment score (Supp. Fig. 8). Once E-Scores for all TFs have been calculated, we fit a linear regression to the distribution of these scores as a function of motif GC-content. Corrected EScores are then calculated from the observed E-Score with the y-offset observed from the linear regression fit (see Eq. 11). This GC bias correction can be optionally turned off.

Subsequently, we assess the significance of the enrichment score by comparison to randomized ROI order, similar to GSEA[56]. To this end, we generate a null distribution of enrichment scores from random permutations, shuffling the rank order of regions and recalculating the E-Score for each shuffled permutation. The final significance of the enrichment score is then calculated from the Z-score, using the Bonferroni correction to account for multiple hypothesis testing. In this manner, TFEA provides a statistically robust and principled way of calculating the motif enrichment that accounts for both differential transcription and motif position in a manner that does not require arbitrary cutoffs.

### 3.4 Differential transcription signal improves motif inference over positional information alone

To assess the effectiveness of the TFEA method, we first compared its performance to both the MD-Score[6] and MDD-Score[52, 24] approaches. We examined a dataset in which a 1 hr Nutlin-3a treatment of HCT116 cells is used to activate TP53[1]. For all methods, sites of RNA polymerase loading and initiation were determined from GRO-seq data[1] using the Tfit algorithm[7] and combined using *muMerge* to identify ROIs. For all methods, the significance cutoff utilized was determined by comparing within treatment replicates (e.g. DMSO to DMSO) and identifying the cutoff at which no changes are detected (see Supp. Fig. 9). Using these per method cutoffs, we recover TP53 from all three approaches (Fig. 3a). Notably, by including differential transcription information, the signal to noise ratio of TP53 detection is significantly improved—modestly in the case of MDD and dramatically for TFEA.

**Figure 3:**
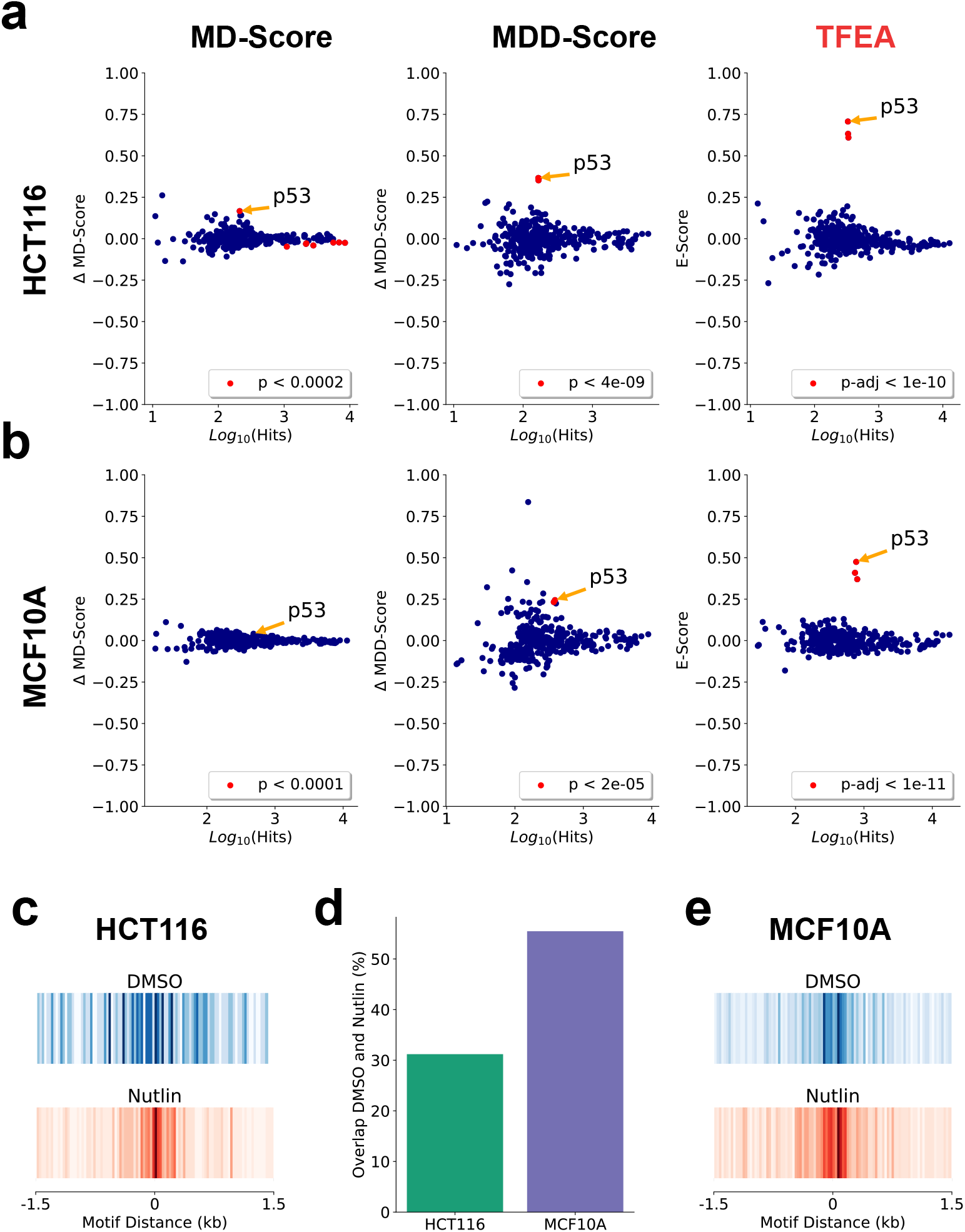
TFEA improves the detection of p53 following Nutlin-3a treatment. (a) Application of the MD-Score, MDD-Score, and TFEA to GRO-Seq data in HCT116 cells with 1hr Nutlin or DMSO treatment [1]. Cutoffs determined by comparing untreated replicates (see Supp Fig 9). (b) Application of the MD-Score, MDD-Score, and TFEA to PRO-Seq data in MCF10A cells with 1hr Nutlin or DMSO treatment. (c) Motif displacement distribution (as heatmap) of TP53 motif instances within 1.5kb of all ROI in either DMSO (blue) or Nutlin-3a (red). (d) Percentage overlap of TP53 motifs within 150bp in DMSO and Nutlin-3a ROIs. (e) Similar to (c) but in MCF10A cells.

We next sought to determine whether TFEA could infer the responsible TF when the underlying changes in transcription were predominantly alterations in existing transcript levels. For this test, we relied on the fact that TP53 response in epithelial cells depends on the TP53 family member TP63[32]. Because TP53 and TP63 have nearly identical motifs, we reasoned that the presence of a constitutively active TP63 would result in elevated basal transcription proximal to TP53/TP63 motifs. To test this hypothesis, we performed PRO-seq on MCF10A cells after 1 hour treatment of either DMSO (control) or Nutlin-3a, and applied all three methods to the resulting data.

Consistent with the constitutive activity of TP63, we observed no change in the TP53 motif by MD-Score analysis (Figure 3b, left). This is due to a larger fraction of ROIs having pre-existing transcription prior to Nutlin-3a exposure in MCF10A relative to HCT116 cells (Figure 3c-e, Supp. Fig. 10). While the MDD-Score method recovers TP53 (Fig. 3b, middle), TFEA significantly improves the signal of the TP53 motif relative to the distribution of all other motifs (Fig. 3b, right). For more detailed analysis of TP53 after Nutlin-3a in HCT116 and MCF10A, see Supp. Figs 11 and 12.

### 3.5 TFEA improves motif enrichment detection by incorporating positional information

We next sought to quantify the performance of TFEA with varying degrees of signal, background, and positional information. As a reference point, we leveraged the widely used MEME-Suite component AME, which quantifies motif enrichment by fitting a linear regression to ranked ROIs as a function of motif instances (Supp. Fig. 13) [45]. To benchmark the two methods, we required biologically representative data sets with known motif enrichment so that error rates could be readily calculated. To this end, we utilized the sites of RNA polymerase initiation detected in untreated GRO-seq datasets of HCT116 cells[1] as the background ROIs. These regions were arbitrarily ordered to mimic a pattern of differential transcription. Subsequently, specific instances of the TP53 motif were generated from the position specific scoring matrix obtained from the HOCOMOCO database[33] and embedded via sequence replacement into the ordered ROI list.

We then varied the number of motifs across ROIs to simulate distinct signal to noise ratios and assess the accuracy of both TFEA and AME (Supp. Fig. 14). Since the significance cutoff thresholds chosen for each method greatly influence the subsequent results, we first measured the mean false positive rate (FPR) and mean true positive rate (TPR) across tests of varying signal and background (Figure 4a). We found that AME detected many false positives (defined as all motifs besides TP53) at loose threshold cutoffs and therefore chose a strict cutoff of 1e-30 for AME. TFEA on the other hand, had a very low FPR even at loose thresholds with the TPR decreasing as the cutoff became stricter. We therefore chose a cutoff of 0.1 for TFEA. We next calculated an F1-Score based on the number of times each method correctly recovered the TP53 motif (and no other motifs) out of the 10 simulations for each test (Fig. 4b).

**Figure 4:**
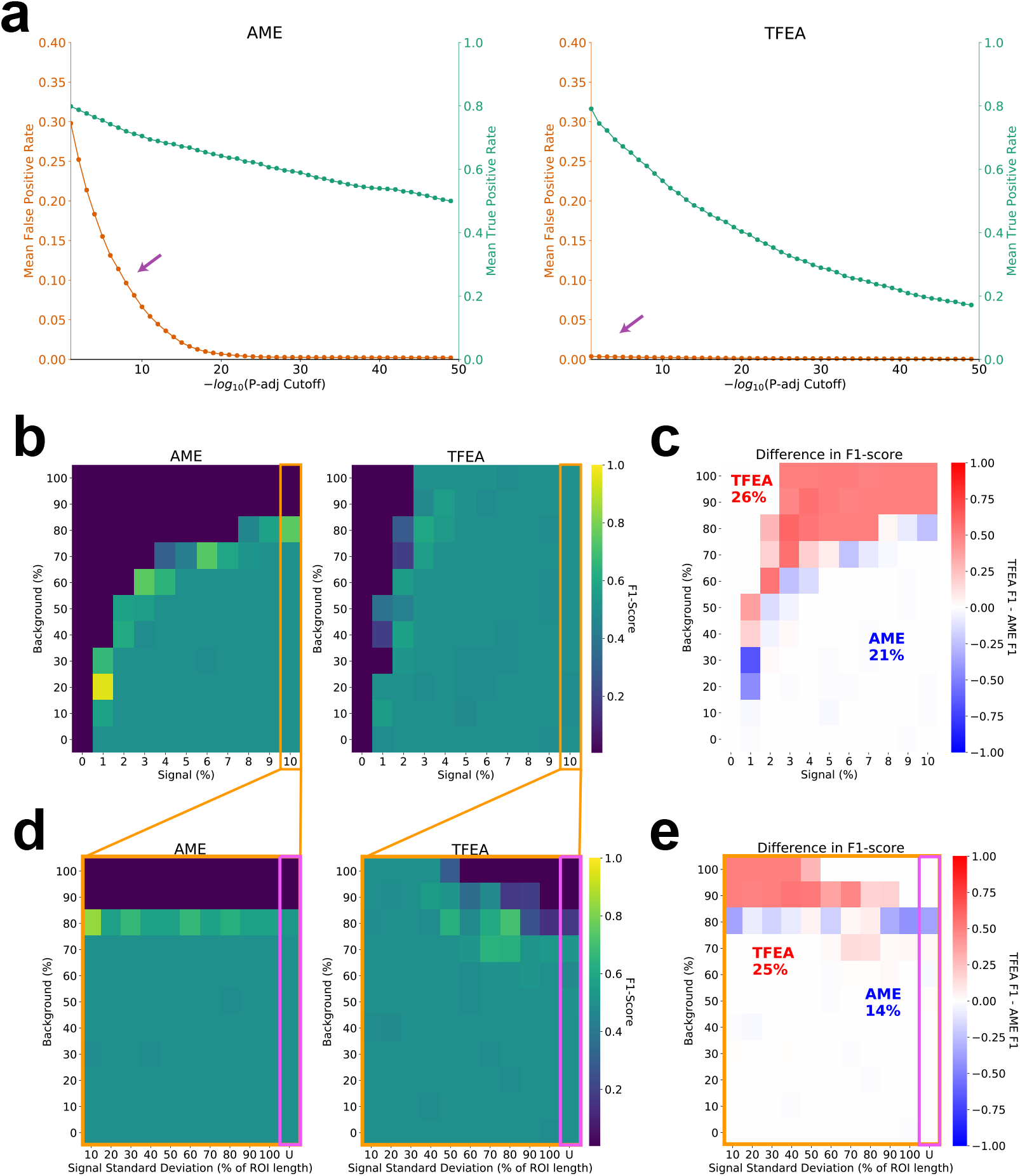
TFEA balances TF positional and differential signal. (a) Optimal cutoffs are determined using the mean true positive rate (TPR; green) and mean false positive rate (FPR; orange) across different signal and background levels as a function of varying the threshold cutoff. (b) F1-score of AME and TFEA for varied signal and background, using optimal AME cutoff 1e-30 and TFEA cutoff 0.1. (c) Difference in F1-score across all simulations (n=121). TFEA outperforms AME in 26% of cases (red) whereas AME outperforms TFEA in 21% of cases (blue). (d) F1-scores and (e) difference in scores for highest signal tested (10% signal), now varying the standard deviation of the signal and background. See Supp Fig 14 for more details on simulations.

We first measured F1-Scores for AME and TFEA with varying relative amounts signal and background (Figure 4b). We found that at high background levels (above 80%), AME was no longer able to detect the enrichment of TP53. TFEA on the other hand, was able to detect TP53 even at high background levels by incorporating positional information. Computing the differential F1-Scores between the two methods (Figure 4c) shows that TFEA performs well in cases where AME detects no enrichment of TP53 (26% of cases), whereas AME outperforms TFEA in 21% of cases.

To further determine how TFEA handles the loss of positional information, we chose the highest signal level tested and altered the variance (standard deviation of the signal position) and the background level (Figure 4d). As expected, AME shows consistent behavior regardless of the positional information of the motif. In contrast, TFEA is able to distinguish signal with differing levels of positional localization. In the extreme case of no positional localization (motifs embedded with a uniform distribution), TFEA performs only slightly worse than AME (Figure 4e).

Additionally, we sought to benchmark the runtime performance and memory usage of TFEA against AME. Here we leverage a first order Markov model (from untreated DMSO samples[1]) to simulate increasing numbers of ROIs as input. Analyzing the core collection of HOCOMOCO TF motifs (n=401), we found that AME runtime increased exponentially while TFEA runtime increased linearly with a single processor (Supp. Fig. 15a). Importantly, TFEA can utilize parallel processing leading to significantly faster runtimes. In terms of memory usage, although TFEA consumes more memory than AME, even in the worst case of 100,000 input regions, TFEA’s memory footprint is less than 1Gb and therefore can still be run on a local desktop computer (Supp. Fig. 15b).

Finally, we sought to examine the performance of TFEA and AME on real data and determine whether TFEA could identify biologically relevant signal in a dataset other than nascent RNA sequencing. Cap analysis of gene expression (CAGE) precisely defines the transcription start site (TSS) of individual transcripts[53, 21, 3]. We analyzed a CAGE-seq timeseries dataset from the FANTOM consortium[21, 9]. In this dataset, human derived monocytes were differentiated into macrophages and treated with lipopolysaccharide (LPS), a proxy for bacterial infection. Differential expression analysis was performed on each LPS time point comparing treatment to control to obtain a list of ranked ROIs.

TFEA recovered the immediate innate immune response, exemplified by the most rapid reported (within 15 min) activation of NF-*κβ* (TF65/RELA, RELB, and NFKB1; Figure 5a). Additionally, TFEA temporally resolved the known secondary response that arises at later time points, which includes the activation of the IFN-stimulated gene factor 3 (ISGF3)[46] complex, comprising IRF9 and STAT1/2[48]. In contrast, AME did not recover the innate immune response at the earliest time point and provided less temporal resolution when distinguishing primary and secondary responses.

**Figure 5:**
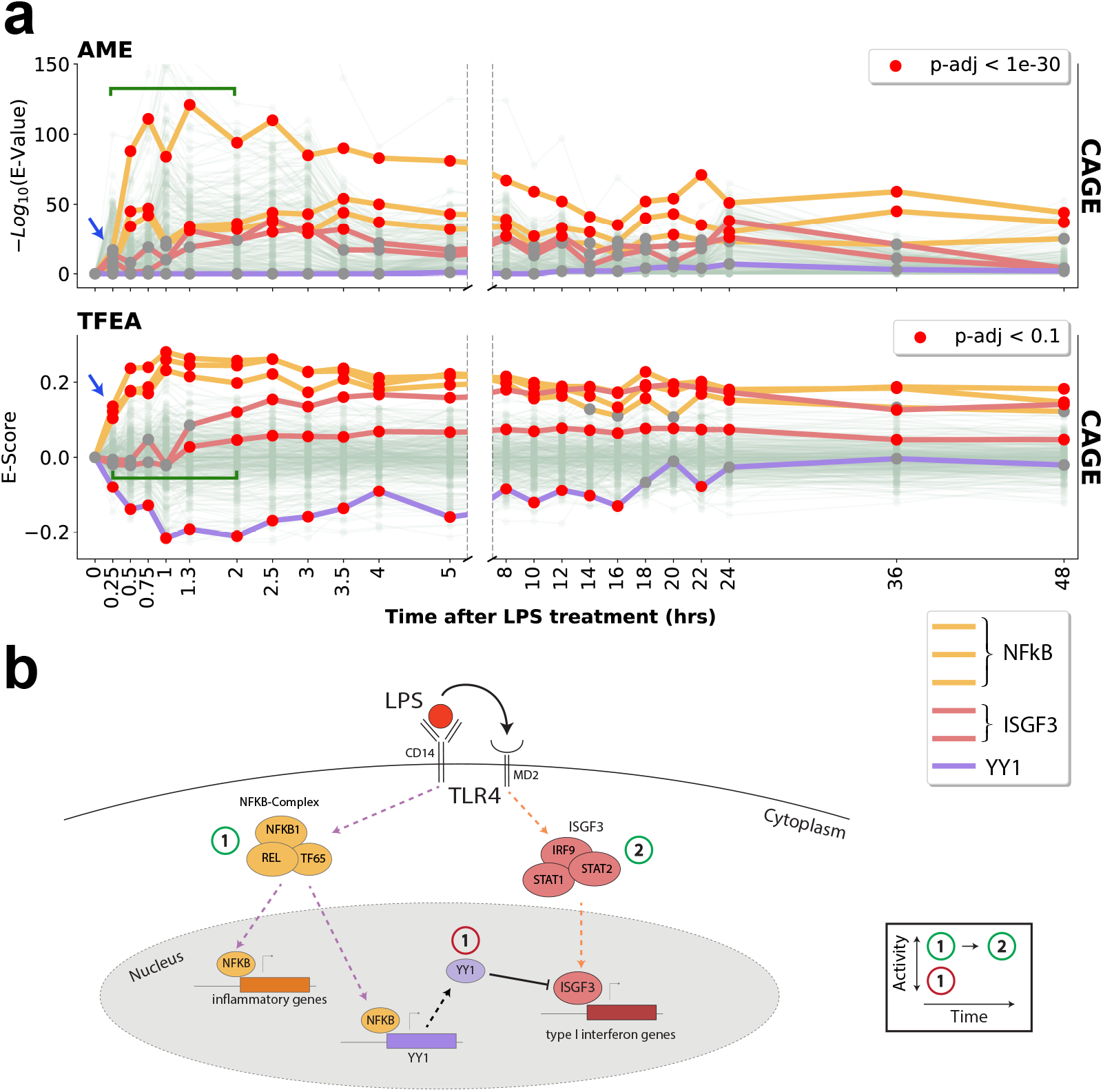
TFEA dissects the temporal dynamics of infection. (a) Analysis of LPS timeseries CAGE data[21, 9] using AME (top) or TFEA (bottom). Trajectories of activity profiles shows LPS triggers immediate activation of the NF-*κβ* complex (TF65/RelB/NFKB1; yellow), observable at 15min (blue arrow). TFEA detects a concomitant down regulation of a set of transcription factors, exemplified here by TYY1 (purple). TFEA also resolves subsequent dynamics (green bracket) of ISGF3 activation (containing IRF9/STAT1/STAT2; red lines). (b) Schematic depicting the molecular insights gained from TFEA analysis. See Supp Fig 16 for more analysis.

Concurrent with the immediate innate immune response, TFEA identified a set of TFs that exhibit a rapid decrease in E-Scores including ELF1/2[17], TYY1 [30][63], USF1/2[31], and GABPA[62]. The decreased E-Score set includes TYY1, a transcriptional inhibitor known to be activated directly by NF*κ*B [54]. Reduction in the E-Score of TYY1 illustrates an important aspect of TFEA— namely, that it cannot distinguish between the activation of a repressor or the loss of an activator. Ultimately, we show with this proof of principle that if the cellular response to LPS was not known *apriori*, we could temporally resolve key aspects of the regulatory network using TFEA and dense time series CAGE data (Figure 5b and Supp. Figure 16).

### 3.6 TFEA works on numerous regulatory data types including ChIP and accessibility data

Though we developed *muMerge* and TFEA for the purpose of inferring TF activity from high resolution data on transcription initiation, this procedure can in principle be used on any assay that produces a localized readout on regulation, such as chromatin immunoprecipitation (ChIP) or DNA accessibility. Although these data sets are less precise and are not direct readouts of polymerase initiation, the popularity of these data make them readily available. To determine whether TFEA could adequately infer TF activity from these datasets, we analyzed a timeseries dataset from ENCODE[19, 44] in which cells were treated with dexamethasone (Dex)—a known activator of the glucocorticoid receptor (GR).

TFEA correctly identifies GR as the key responding TF from the datasets that most closely capture RNA polymerase initiation (including p300, H3K27ac, and DNA accessibility), and does not identify GR for the transcriptionally repressive mark H3K9me3 (Figure 6a)[40, 44]. Surprisingly, the effects of p300 and H3K27ac are seen rapidly, as soon as 5min after dexamethasone treatment. As expected, H3K27ac deposition is temporally lagged behind its canonical acetyl-transferase p300[29, 60, 51]. Additionally, the enhancer marks H3K4me1 and H3K4me2 show strong enrichment of GR by 30min but the promoter mark H3K4me3 shows only modest enrichment, further supporting the finding that GR binds primarily at enhancers[44] (Supp. Fig. 17). Using the diversity of data types and dense time series, we can construct a temporally resolved mechanism of how GR effects changes in transcription (Figure 6b and c). In short, TFEA’s results for this array of accessibility marks are exactly consistent with biological expectation.

**Figure 6:**
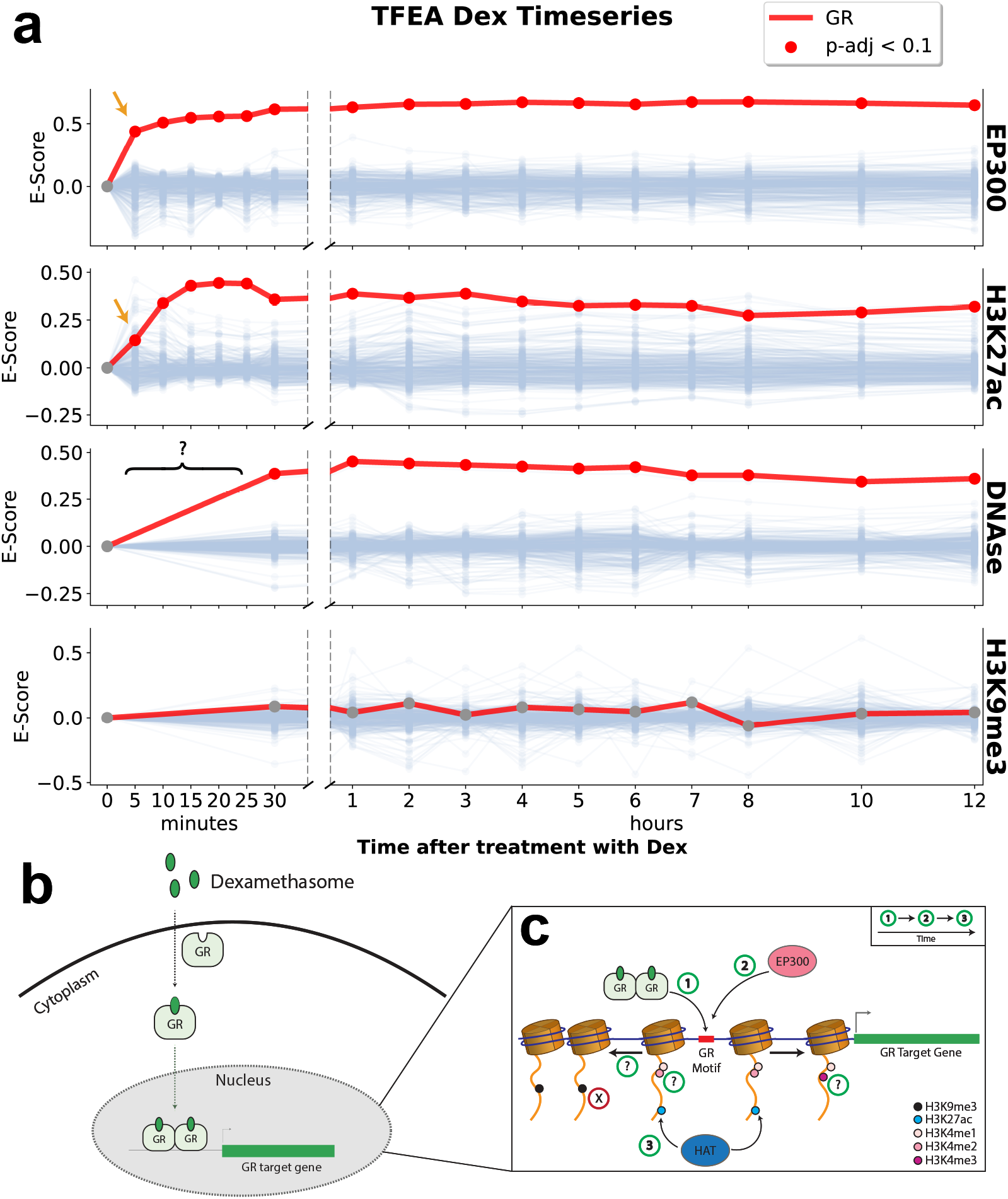
TFEA captures rapid dynamics of glucocorticoid receptor (GR) following treatment with dexamethasone. (a) TFEA correctly identifies GR from time series ChIP data on the histone acetyl-transferase p300, H3K27ac and DNase I. No signal is observed in the negative control H3K9me3. TFEA correctly shows a temporal lag in H3K27ac signal (yellow arrow). (b) Known cellular dynamics of GR induced by dexamethasone. (c) Mechanistic and temporal insights gained by performing TFEA analysis, question marks indicate datasets where earlier time points were not available to resolve temporal information.

### 3.7 Discussion

We present here transcription factor enrichment analysis (TFEA), a computational method that measures the global correlation between the position of a TF motif and its differential effects on transcription across the genome, following any given perturbation. We show that TFEA outperforms existing enrichment methods when positional data is available and is comparable to these methods in the absence of positional signal. Further, we show that TFEA, when leveraged with high resolution time series data, can provide mechanistic insight into the order of regulatory events responding to the perturbation.

A key aspect of TFEA is the incorporation of both positional and differential information in calculating TF activity. Most current motif enrichment algorithms use solely differential information, likely due to the poor positional resolution on historically popular techniques such as ChIP-Seq. Methods such as nascent transcription and CAGE provide higher resolution on the position of RNA polymerase initiation genome wide. To leverage the improved resolution of these methods, we introduce *muMerge*, a statistically principled way of combining ROIs across replicates and conditions that better captures position and lengthscale information as compared to standard merging or intersecting approaches. The presence of improved positional information greatly increases the ability to detect biologically relevant TFs.

Although TFEA makes significant improvements in detecting the activity of TFs in response to perturbations, there are several aspects of this approach that could be improved. TFEA is dependent on having a collection of known motifs, yet some TFs have no known motif or one of poor quality. However, over time, the quality and numbers of TFs in the major databases have dramatically improved[38]. Furthermore, TFEA can only distinguish between paralogous motifs to the extent that they have distinct motifs. Importantly, motif scanning still requires a fixed cutoff within TFEA. Future iterations of the method could conceivably eliminate this cutoff, but likely this will substantially increase runtimes for what may only be minor gains in performance. Genome-wide, sites of transcription initiation (both promoters and enhancers) show substantial GC bias. Often short high GC content motifs, which are exceedingly common in ROIs, appear to show significant changes with a perturbation. While we made some effort to account for this using linear regression, this approach is empirical and a more principled approach is desired.

Despite these caveats, TFEA recovers known TF dynamics across a broad range of data types in response to a variety of perturbations. Inevitably, the data type utilized influences the detection ability of TFEA. For example, while CAGE data provides precise resolution on the TSS, it must be deeply sequenced to reliably detect enhancer associate transcription events[15]. Consequently, TFs that predominantly regulate enhancers will likely be less detectable in poorly sequenced CAGE data. On the other hand, some methods are more capable of detecting immediate changes in RNA polymerase initiation, allowing for shorter more refined time points. As demonstrated here, TFEA is able to leverage the information from each data set by incorporating both its distinct positional and differential signal. Applying TFEA to diverse data types, using dense time series, can uncover a detailed mechanistic understanding of the key regulators that enact the cell’s dynamic response to a perturbation.

## 4 Online Methods

### 4.1 TFEA

We have developed Transcription Factor Enrichment Analysis (TFEA) to identify transcription factors that demonstrate significant differential activity following a perturbation. It has been observed that, during a perturbation, the binding sites of active transcription factors co-localize with regulatory regions that exhibit strong differential RNA polymerase initiation[6]. TFEA leverages this observation to calculate an enrichment score that quantifies this activity and an associated significance for each TF.

Here we describe in detail the key steps of the TFEA pipeline (shown in Figure 1)—specifically, for each TF we describe how the main input (regions of interest—ROIs) are defined, how the ROIs are ranked, and how the enrichment score is subsequently calculated and GC-corrected.

#### 4.1.1 Defining the Regions of Interest with *muMerge*

One input required for TFEA is a common set of regions of interest (ROIs) on which all experimental samples are evaluated. Each region (consisting of a genomic start and stop coordinate) represents a reference point (the midpoint of the region) and an uncertainty on that reference point (the width of the region). Biologically, the reference point is the presumed transcription start site. Regions can be derived from a number of data types, with varying degrees of precision. For example, CAGE data provides a highly precise measure of a TSS while nascent sequencing is slightly less precise. Other assays like ChIP (for RNA polymerase or H3K4 methylation) or ATAC have much lower positional precision.

Regardless of the assay, most methods for identifying such regions fit each dataset independently (*e.g*., a peak caller for ChIP data or Tfit for identifying sites of bidirectional transcription in nascent data). As a result, these regions will not be exactly consistent between samples (*e.g*. some sites are condition specific and even for shared sites boundaries may vary). Therefore, a method is needed to combine the regions from all the samples into a consensus set. To this end, we developed a probabilistic, principled method (hereafter referred to as *muMerge*) for determining consensus regions of interest, informed by the corresponding regions predicted from individual samples. *muMerge* was developed specifically for determining the set of consensus RNA polymerase loading and initiation sites observed in nascent sequencing data (by combining bidirectional calls from any number of samples) but it can also be applied to peak calls generated from numerous other regulatory data types (*e.g*., ChIP, ATAC, or histone marks).

The basic assumption made by *muMerge* is that each sample is an independent observation of an underlying set of hypothetical loci—where each hypothetical loci has a precise critical point *μ*, of which the corresponding sample region ([*start*, *stop*]) is an estimate. We assume this loci is more likely to be located at the center of the sample region than at the edges, so *muMerge* represents the sample region by a standard normal probability distribution, centered on the region, whose standard deviation correlates with region width.

To calculate a best estimate (the ROI) for a given loci, *muMerge* calculates a joint probability distribution across all samples from all regions that are in the vicinity of the loci. This joint distribution is calculated by assuming:

1. replicates within a condition are independent and identically distributed (*i.i.d*.)
2. replicates *across* conditions are mutually exclusive (*i.e*., a sample cannot represent multiple experimental conditions)

Hence *muMerge* computes the product of the normal distributions across all *replicates* within a condition and then sums these results across all *conditions*. The best estimates for the transcription loci *μ* are taken to be the maxima of this joint distribution—these are the ROI positions. Finally, to determine an updated width, or confidence interval, for each ROI, *muMerge* assumes that the original sample regions whose midpoints are closest to the new position estimate are the most informative for the updated width. Thus the ROI width is calculated by a weighted sum of the widths of the original regions, weighted by the inverse of the distance to each one.

##### *muMerge* mathematical description

Principally, *muMerge* makes two probabilistic assumptions about sequenced samples:

- **Assumption A:** Replicate samples are independent measurements of *identical experimental conditions* and therefore any corresponding sample regions within them are independent and identically distributed (*i.i.d*.) observations of a common random variable (*i.e*., the underlying hypothetical loci).
- **Assumption B:** Cross-condition samples are independent measurements of *mutually exclusive experimental conditions* and therefore any sample regions within them are observations of (potentially) disjoint random variables.

These two assumptions inform how *muMerge* accounts for each individual sample, when computing the most likely ROI for any given genomic location (see below for further details).

To start, the two inputs to *muMerge* are a set of regions for each sample (genomic coordinates: {[*start*, *stop*],…}) that annotate the sequenced features present in the dataset, as well as an experimental conditions table that indicates the sample groupings (which samples are from which experimental condition). With these inputs, *muMerge* performs the following steps to compute a global set of ROIs:

1. Group overlapping sample regions, processing each group one at a time
2. Express each sample region as a positional probability distribution
3. Generate a joint distribution
4. Identify maximum likelihood ROI positions from the joint distribution
5. Compute ROI widths via weighted sum
6. Adjust the sizes of overlapping ROIs
7. Record final ROIs for the given group
8. Repeat 2–8 for all remaining groups

First, from the input samples, *muMerge* groups all sample regions that overlap in genomic coordinate (a region is grouped with all other regions it overlaps and, transitively, with any regions overlapping those). We denote a single group of overlapping regions as *G_r_*. This grouping is done globally for all samples, resulting in a set of grouped regions *G* = *G_r_*, such that every sample region is contained in exactly one grouping *G_r_* (i.e., 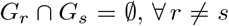). Then each group of regions, *G_r_*, is processed individually, as the remainder of this section describes. For a given group, we denote each sample region within it as the 2-tuple (*μ_k_*, *σ_k_*)_*ij*_ ∈ *G_r_*, where *μ_k_* is the genomic coordinate (base position) of the center of the region and *σ_k_* is the region half-width (number of bases). The indices denote the *k*-th sample region for replicate *j* in condition *i*.

*muMerge* then processes the regions in *G_r_* as follows. Each region within the group is expressed as a standard normal distribution (*ϕ*) as a function of base position *x*,

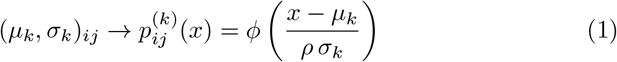

where *ρ* is the “width ratio”— the ratio of the half-width sample region to the standard deviation of the normal distribution—with a default of *ρ* = 1 (user option). This distribution represents the probability of the location for the underlying hypothetical loci (*μ*), of which (*μ_k_*, *σ_k_*)_*ij*_ is an estimate. For those samples with no regions within *G_r_*, the probability distribution is expressed as a uniform, 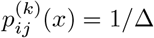 where Δ is the full range encompassed by the overlapping sample regions. In other words, we assume that if the sample contains no data to inform the location of the underlying loci at that location, then all positions are equally likely for that sample. *muMerge* then calculates a joint distribution 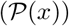 by combining all 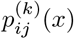 for the group as follows:

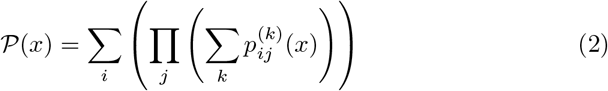

Here we are calculating the product of the replicate distributions (index *j*—those within a given experimental condition), consistent with our probabilistic assumption A, and the sum of the resulting distributions across experimental conditions (*i* index), consistent with our probabilistic assumption B. Though this function is not a normalized probability distribution, we are only interested in relative values of 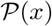. Specifically, we are interested in the maxima of this function. We identify the set of maxima (which we denote 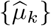) and rank them by the function value for each position, 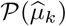. We then keep the top *M* + 1 from the ranked set, where *M* is the median number of regions per sample in *G_r_* (user option). This is our final set of estimates on the hypothetical loci positions, *μ*—*i.e*., the positions of our ROIs for group *G_r_*.

For each 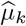, we then calculate a width for the resulting ROI. We do so for each by calculating a weighted sum over the set of all original sample regions in the group, {(*μ_k_*, *σ_k_*)_*ij*_}, weighted by the inverse of the distance from the final position estimate to each *μ_k_*. Thus the final ROI half-width, 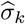, is calculated as follows:

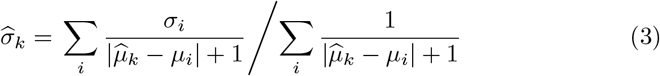

where *i* indexes all sample regions in the group *G_r_* = {(*μ_k_*, *σ_k_*)_*ij*_}. Our rationale is that the width of those sample regions that are closer to the ROI position 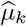, are more informative for the ROI width and therefore are given a larger weight. This results in a set of ROIs 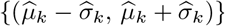.

Finally, we determine if there is overlap between any of the regions in this set of ROIs. If so, any two overlapping regions are reduced in size, symmetrically about their centers, until they no longer overlap. This is done so that any genomic position can be uniquely associated with an ROI. The final ROIs for the group are then written to an output file to be used downstream in the pipeline. This process is repeated for all groups of overlapping sample regions (i.e., ∀ *G_r_* ∈ *G*).

#### 4.1.2 Ranking ROIs

With a set of ROIs identified, the next step is to rank them by differential signal. Because the goal of TFEA is to identify transcription factors that are enriched during a perturbation and because the ROIs are associated with transcription factor activity, it follows that a ranking based on the differential signal at the ROIs would capture the regulatory behavior of the TF. For different types of datasets, the differential signal represents different biological processes—differential transcription for nascent (PRO-seq or GRO-seq), differential accessibility (DNAse or ATAC-Seq), and differential occupancy for ChIP. There are a number of ranking metrics one could use that are based on these differential signals—for example, difference in coverage, log-fold change, or a differential significance (p-value). For TFEA, we chose to rely on a well-established tool (*DESeq2*) to perform our ranking, since it was designed to model the statistical variation found in sequencing data[41].

For a set of ROIs, TFEA calculates read coverage for each replicate and condition using *bedtools multibamcov* (version 2.25.0)[50]. TFEA then inputs the generated counts table into *DESeq2* [41] (or *DESeq* [2] if no replicates are provided) to obtain differential read coverage for all ROIs. By default, these regions are then ranked by the *DESeq2* computed p-value, separated by positive or negative log-fold change (alternative user option to rank the ROIs by foldchange). In other words, the ROIs are ranked from the most significant positive fold-change to the most significant negative fold-change.

#### 4.1.3 Identifying location of motif instances

Accurately identifying the locations of motif instances relative to each ROI is a critical step in the TFEA pipeline. By default TFEA uses the motif scanning method FIMO, which is a part of the MEME suite (version 5.0.3)[23]. FIMO represents each TF by a base-frequency matrix and uses a zero-order background model to score each position of the input sequences. For each ROI, we scan the 3kb sequence surrounding the ROI center 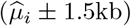. This 3kb window was chosen primarily to reduce computation time and is also consistent with the window used for the MD-score method[6]. For each TF, we utilize a scoring threshold of 10^-6^ and keep the highest scoring position (denoted *m_i_*), in the event more than one motif instance is identified. If no position score above the threshold, then no *m_i_* is recorded for the ROI. Our background model is determined by calculating the average base frequency over all 3kb regions. For each TF, we use the frequency-matrix from the HOCOMOCO database[33] with a default psuedo-count of 0.1.

#### 4.1.4 Enrichment Score

With the motif instances identified for each of the ranked ROIs, we now detail how TFEA calculates the enrichment score (“E-Score”—in Fig. 1) for each transcription factor. The procedure for calculating enrichment requires two inputs:

1. N-tuple sequence 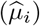—the genomic coordinates for reference points, assumed to be the centers of all ROIs (*e.g*., consensus ROIs calculated by *muMerge*), ranked by *DESeq2* p-value (separated by the sign of the fold-change).
2. Sequence (*m_i_*)—the genomic coordinates of each max-scoring motif instance (*e.g*., motif loci generated by scanning with FIMO), for each ROI.

We first calculate the motif distance *d_i_* for each ROI—the distance from each 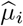 to the highest scoring motif instance *m_i_* within 1.5kb of 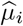. If no *m_i_* exists within 1.5kb, then *d_i_* is assigned a null value (Ø) (Eq. 4).

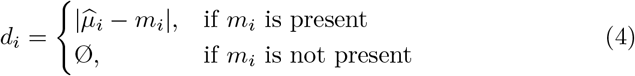

Next, we calculate the background distribution of motif distances. We assume the majority of the ROIs experience no significant fold-change—namely, those ROIs in the middle of the ranked list. Consequently, we calculate the mean, background motif distance (Eq. 5) for those ROIs whose rank is between the first and third quartiles of the sequence of ROI positions, 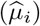, as follows

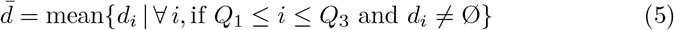

where *Q*_1_ and *Q*_3_ are the first and third quartiles, respectively. Our assumption is that the inter-quartile range of the sequence 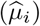—between indices *Q*_1_ and *Q*_3_—represents the background distribution of motif distances for the given transcription factor, and therefore defines the weighting scale for significant ROIs in our enrichment calculation. We found this to be essential since the background distribution varies between transcription factors. This variation in the background can be attributed to the similarity of a given motif to the base content surrounding the center of ROIs. For example, in the case of RNA polymerase loading regions identified in nascent transcription data (which demonstrate a greater GC-content proximal to *μ* as compared to genomic background[6]), GC-rich transcription factor motifs were more likely to be found proximal to each ROI by chance and thus resulted in a smaller 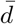 than would be the case for a non-GC-rich motif.

Having calculated the mean background motif distance, we proceed to calculate the enrichment contribution (*i.e*., weight—Eq. 6) for each ROI in the sequence (see “Weight Calculation” in Fig. 1).

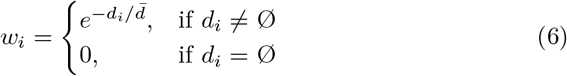

In order to calculate the E-Score, we first generate the enrichment curve for the given TF (solid line in “Enrichment Curve” in Fig. 1) and the background (uniform) enrichment curve (dashed line in “Enrichment Curve” in Fig. 1). We define the E-Score as the integrated difference between these two (scaled by a factor of 2, for the purpose of normalization). The enrichment curve (Eq. 7), which is the normalized running sum of the ROI weights, and the E-Score (Eq. 8) are calculated as follows:

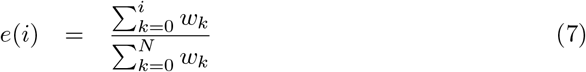

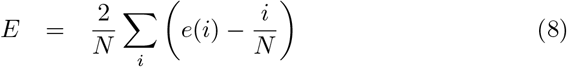

where *i* is the index for the ROI rank and *i*/*N* represents the uniform, background enrichment value for the *i*th of *N* ROIs. The background enrichment assumes every ROI contributes an equal weight *w_i_*, regardless of its ranking position. Therefore, the enrichment curve (Eq. 7) will deviate significantly from background if there is correlation between the weight and ranked position of the ROIs. In this case, the E-Score will significantly deviate from zero, with *E* > 0 indicating either increased activity of an activator TF or decreased activity of a repressor TF. Likewise, *E* < 0 indicates either a decrease in an activator TF or an increase in a repressor TF. By definition, the range of the E-Score is −1 to +1.

Unlike GSEA, which uses a Kolmogorov–Smirnov-like statistic to calculate its enrichment score[56], the TFEA E-Score is an area-based statistic. GSEA was designed to identify if a predetermined, biologically related subset of genes is over-represented at the extremes of a ranked gene list. Therefore, the KS-like statistic is a logical choice for measuring how closely clustered are the elements of the subset, since it directly measures the point of greatest clustering and otherwise is insensitive to the ordering of the remaining elements. Conversely, because TFEA’s ranked list does not contain two categories of elements (the ROIs) and all elements can contribute to the E-Score, we wanted a statistic that was sensitive to how all ROI in the list were ranked—for this reason, we chose the area-based statistic. The null hypothesis for TFEA assumes all ROI contribute equally to enrichment, regardless of their motif co-localization and rank. Hence the uniform background curve, to which the enrichment curve is compared.

In order to determine if the calculated E-Score (Eq. 8) for a given transcription factor is significant, we generate a E-Score null distribution from random permutations of 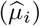. We generate a set of 1000 null E-Scores 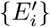, each calculated from an independent random permutation of the ranked ROIs, 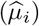. Our E-Score statistic is zero-centered and symmetric, therefore we assume 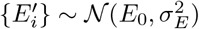. The final E-Score for the transcription factor is compared to this null distribution to determine the significance of the enrichment.

Prior to calculating the E-Score p-value, we apply a correction to the E-Score based on the GC-content of the motif relative to that of all other motifs to be tested (user configurable). This correction was derived based on the observation that motifs at the extremes of the GC-content spectra were more likely to called as significant across a variety of perturbations. We calculate the E-Scores for the full set of transcription factors as well as the GC-content of each motif, {(*g_i_*, *E_i_*)}. We then calculate a simple linear regression for the relationship between the two,

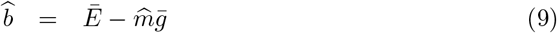

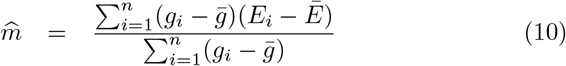

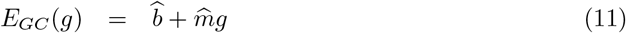

where 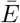 and 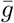 are the average E-Score and average GC-content. *E_GC_*(*g*) is the amount of the E-Score attributed to the GC-bias for a motif with GC-content *g*. Thus the final E-Score for the transcription factor is given by *E_TF_* = *E* − *E_GC_*(*g_TF_*), the difference between Eq. 8 and 11. If GC-content correction is not performed, then Eq. 8 is taken to be the final E-Score. The p-value for the final TF E-Score is then calculated from the Z-score, *Z_TF_* = (*E_TF_* − *E*_0_)/*σ_E_*.

### 4.2 Software Availability

TFEA is available for download at https://github.com/Dowell-Lab/TFEA and comes with muMerge integrated. Alternatively, *muMerge* can be downloaded independently at https://github.com/Dowell-Lab/mumerge. Additionally, TFEA can be utilized through the web interface at https://tfea.colorado.edu/.

### 4.3 Benchmarking

In order to benchmark the performance of *muMerge* and TFEA, we performed a number of simulations that isolate the different parameters of *muMerge* and TFEA, comparing the performance to that of some commonly used alternatives. Here we describe how the data for each test was generated.

#### 4.3.1 *muMerge*: Simulating replicates for calculation of ROIs

To test the performance of *muMerge* in a principled manner, we first generate replicate data in a way that simulates the uncertainty present in individual samples. For each replicate, we perform 10,000 simulations of sample regions for a single loci, and calculate the average performance. For each simulation we assume a precise position and width for the hypothetical loci and model the uncertainty of each sample region with a binomial and Poisson distribution, respectively. The position of each sample region, 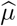, is pulled from a symmetric binomial distribution 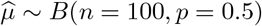, centered at zero. The half-width of each sample region, *W*, is pulled from a Poisson distribution *W* ~ *Pois*(λ = 100). The specific distributions utilized to generate the sample regions are as follows:

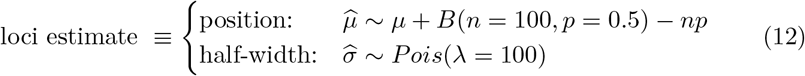

Here *B*(·) is the binomial distribution centered at *np* with success probability 0.5 and variance *np*(1 − *p*) = 25. Thus, the position estimator 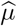 is centered at *μ*. *Pois*(·) is the Poisson distribution, thus, the half-width for each sample region have mean and variance of λ = 100.

The first test consisted of inferring a single loci (located at *μ* = 0) from an increasing number of replicates. A sample region for each replicate was generated from Eq. 12. This simulation was repeated 10,000 times for each number of replicates being combined. The methods *muMerge*, *bedtools merge* and *bedtools intersect* were applied to each of the 10,000 simulations. The average error on the midpoint (its deviation from the true loci position, *μ* = 0) and region width were calculated for the regions generated from each method, averaged over all 10,000 simulations. The behavior of the average positional error and region width as a function of number of combined replicates is shown in Fig. 2a, b.

The second test consisted of inferring two loci (*μ*_1_ = −*x* and *μ*_2_ = +*x*) as the distance between those loci was increased (from *x* = 0 to 200). This simulation was repeated 10,000times for each value of *x* (with 3 replicates). The distribution of the inferred positions and widths were plotted, using *muMerge*, *bedtools merge* and *bedtools intersect*. The distribution of positions and widths as a function of the distance between *μ*_1_ and *μ*_2_ are shown in Fig. 2c, d.

#### 4.3.2 TFEA: Simulated motif enrichment

To generate test sequences for benchmarking, we randomly sampled 10,000 sequences from detected bidirectionals in untreated HCT116 cells [1]. We then embedded instances of the TP53 motif in the highest ranked sequences with a normal distribution with *μ* = 0 and *σ* = 150 (representative of signal). To simulate background noise, we embedded instances of the TP53 motif with a uniform distribution to a percentage of the remaining sequences (chosen randomly). To calculate an F1-Score, for each scenario of varying signal to background we generated 10 simulations. We then calculated the harmonic mean of precision and recall with the aggregate p-values of all 10 simulations measuring all 401 TF motifs within the HOCOMOCO database (total 4010 TF motifs). True positives, in this case, were the 10 instances of the TP53 motif that should be significantly enriched. Any other significantly enriched TF motifs were considered false positives. We performed two sets of tests: 1) varying the amount of signal and the amount of background and 2) varying the standard deviation of the highest signal tested (10% signal; with the last scenario being uniform signal distribution) and the amount of background.

#### 4.3.3 TFEA: Testing compute performance

The base (ATGC) content of regulatory regions was calculated from the sites of RNA polymerase initiation inferred in HCT116 DMSO (using Tfit; described in [6]). One million 3kb sequences were generated based on the empirical probability of the positional base composition. We then randomly sampled an increasing number of sequences (up to 100,000) to be used in the computational processing tests. Run time and compute resources were measured using the Linux *time* command on a single node of a 70-node mixed-platform high-memory compute cluster running CentOS 7.4. To compute the runtime for a single processor, we added the *systime* and *usertime*. To compute memory usage for a single processor, we reran TFEA using only a single processor.

### 4.4 Datasets Utilized

We generated PRO-seq libraries for MCF10A cells with and without Nutlin-3a. Additionally, a number of publicly available datasets were utilized, including: Allen 2014 (Nutlin-3a, GC-correction), ENCODE (GGR: Reddy - Dex/GR) and FANTOM (Baillie - Macrophage/LPS). See supplemental material for a full list of accession codes.

#### 4.4.1 PRO-Seq in MCF10A

##### Cas9RNP formation

sgRNA was formed by adding tracrRNA (IDT cat# 1072533) and crRNA (TP53 exon 2, positive strand, AGG PAM site, sequence: GATCCACTCACAGTTTCCAT) in a 1:1 molecular ratio together and then heating to 95°C and then allowing to slowly cool to room temperature over 1 hour. Cas9RNP was then formed by adding purified Cas9 protein to sgRNA at a ratio of 1:1.2. 3.7*μ*L of purified Cas9 protein at 32.4*μ*M was added to 2.9*μ*L of 50*μ*M sgRNA. This was then incubated at 37°C for 15 minutes, and used at 10*μ*M concentration within the hour.

##### Donor Plasmid Construction

Vector Builder was used to construct plasmid. Insert was flanked by 1.5kb homology arms, and mCherry was inserted as a selection marker.

##### CRISPR/Cas9 Genome Editing

MCF10A cells cultured in DMEM/F12 (Invitrogen #11330-032) media containing 5% horse serum (LifeTech #16050-122), 20ng/mL EGF ((Peprotech #AF-100-15), 0.5*μ*g/mL Hydrocortisone (Sigma #H0888-1g), 100ng/mL Cholera toxin (Sigma #C8052-2mg), 10*μ*g/mL insulin (Sigma #I1882-200mg), and 1x Gibco 100x Antibiotic-Antimycotic (Fisher Sci, 15240062) penicillin-streptomycin. Cells were split 24 hours prior to experiment and grown to approximately 70% confluency on a 15cm plate. Media was aspirated, and the cells were washed with PBS. 4ml of trypsin per plate were used to harvest adherent cells, after which 8mL of resuspension medium (DMEM/F12 containing 20% horse serum and 1x pen/strep) was added to each plate to neutralize the trypsin. Cells were collected in a 15ml centrifuge tube and spun down at 1,000xg for 5 minutes, then washed in PBS and spun down again at 1,000xg for 5 minutes. Cells were counted using a hemocytometer and 5*x*10^5^ cells were put in individual 1.5mL eppedorph tubes for transfection. Cells were re-suspended in 4.15*μ*L Buffer R, 10*μ*M Cas9RNP (6.6*μ*L), 1*μ*g WTp53 donor plasmid (1.25*μ*L). Mixture was drawn up into a 10*μ*L Neon pipet tip, electroporated using the Neon Transfection Kit with 10*μ*L tips (1400V, 20ms width, 2 pulse). Transfected cells were then pipetted into 2mL of antibiotic free media. After 1 week of recovery, cells were then single cell sorted into 96 well plate based on mCherry expression. Clones were then verified with sequencing, PCR, and western blot.

##### Nuclei Preparation

MCF10A WTp53 cells were seeded on three 25cm dishes (1×107 cells per dish) for each treatment 24 hours prior to the experiments (70% confluency at the time of the experiment). Cells were treated simultaneously with 10*μ*M Nutlin3a or 0.1% DMSO for 1 hour. After treatment, cells were washed 3x with ice cold PBS, and then treated with 10 ml (per 15 cm plate) ice-cold lysis buffer (10 mM Tris–HCl pH 7.4, 2 mM MgCl2, 3 mM CaCl2, 0.5% NP-40, 10% glycerol, 1 mM DTT, 1x Protease Inhibitors (1mM Benzamidine (Sigma B6506-100G), 1mM Sodium Metabisulfite (Sigma 255556-100G), 0.25mM Phenylmethylsulfonyl Fluoride (American Bioanalytical AB01620), and 4U/mL SUPERase-In). Cells were centrifuged with a fixed-angle rotor at 1000xg for 15 min at 4°C. Supernatant was removed and pellet was resuspended in 1.5 mL lysis buffer to a homogenous mixture by pipetting 20-30X before adding another 8.5 mL lysis buffer. Suspension was centrifuged with a fixed-angle rotor at 1000xg for 15 min at 4°C. Supernatant was removed and pellet was resuspended in 1 mL of lysis buffer and transferred to a 1.7 mL pre-lubricated tube (Costar cat. No. 3207). Suspensions were then pelleted in a microcentrifuge at 1000xg for 5 min at 4°C. Next, supernatant was removed and pellets were resuspended in 500 *μ*L of freezing buffer (50 mM Tris pH 8.3, 40% glycerol, 5 mM MgCl2, 0.1 mM EDTA, 4U/ml SUPERase-In). Nuclei were centrifuged 2000xg for 2 min at 4°C. Pellets were resuspended in 100 *μ*L freezing buffer. To determine concentration, nuclei were counted from 1 *μ*L of suspension and freezing buffer was added to generate 100 μL aliquots of 10*x*10^6^ nuclei. Aliquots were flash frozen in liquid nitrogen and stored at −80°C.

##### Nuclear run-on and RNA preparation

Nuclear run-on experiments were performed as described (Mahat et al., 2016) with the following modifications: the final concentration of non-biotinylated CTP was raised from 0.25 *μ*M to 25 *μ*M, a clean-up and size selection was performed using 1X AMPure XP beads (1:1 ratio) (Beckman) prior to test PCR and final PCR amplification, and the final library clean-up and size selection was accomplished using 1X AMPure XP beads (1:1 ratio) (Beckman).

##### Sequencing

Sequencing of PRO-Seq libraries was performed at the BioFrontiers Sequencing Facility (UC-Boulder). Single-end fragment libraries (75 bp) were sequenced on the Illumina NextSeq 500 platform (RTA version: 2.4.11, Instrument ID: NB501447), demultiplexed and converted BCL to fastq format using bcl2fastq (bcl2fastq v2.20.0.422); sequencing data quality was assessed using FASTQC (v0.11.5) and FastQ Screen (v0.11.0), both obtained from https://www.bioinformatics.babraham.ac.uk/projects/. Trimming and filtering of low-quality reads was performed using BBDUK from BBTools (v37.99) (Bushnell, n.d.) and FASTQ-MCF from EAUtils (v1.05) [5].

##### Availability

MCF10A PRO-seq data is available in GEO with accession numbers GSE142419.

#### 4.4.2 Data Processing

##### GRO/PRO-Seq data

All GRO-Seq and PRO-Seq data were processed using the Nextflow[20] NascentFlow pipeline (v1.1 [59]) specifying the ‘–tfit’ flag. Subsequent Tfit bed files from all samples were combined with *muMerge* to obtain a consensus list of ROIs.

##### ENCODE data

Raw bed and bam files were downloaded directly from ENCODE (encodeproject.org). These files were inputted directly into the TFEA pipeline for processing and analysis. AME analysis was performed on the ranked ROI list produced as an optional output from TFEA.

##### FANTOM data

Raw expression tables for the Macrophage LPS time series were downloaded using the table extraction tool (TET) from the FANTOM Semantic catalogue of Samples, Transcription initiation, And Regulations (SSTAR; http://fantom.gsc.riken.jp/5/sstar/Macrophage_response_to_LPS). Because the annotations for regions within hg38 counts tables contained “hg19”, we performed this analysis in the hg19 genome with the hg19 counts table instead of the hg38 counts table. We then performed DE-Seq analysis (since there were no replicates) on each time point compared to control and ranked the annotated regions within the counts table similar to Figure 1. We then ran TFEA and AME with default settings on each of the three donors. We displayed only data for donor 2, as this sample had the most complete time series data.

##### Clustering FANTOM data

We retained TFs with at least 15 significant (p-adj < 0.1) time points (representing 2/3 of all timepoints) from the TFEA output and applied K-means clustering. Clustering of the time series data was performed on the first two hours only, in order to distinguish the early responses to LPS infection. K-means clustering was conducted using the Hartigan and Wong algorithm with 25 random starts and 10 iterations for *k* = 3 [27]. The optimal number of clusters was selected using the Elbow method [13].

##### String database analysis

Protein names from TFs that were found to be significant in at least 15 time points were taken from the HOCOMOCO database. These proteins were inputted directly into the String database (https://string-db.org). Clusters were formed by selecting the MCL clustering option with an inflation parameter of 3 (default). Network edges were selected to indicate the strength of the data support. Finally, nodes disconnected from the network were hidden.

## Supporting information

All Supplemental Material

## Acknowledgments

This work was funded in part by a National Science Foundation (NSF) ABI grant number 1759949, a National Institutes of Health (NIH) grant RO1 GM125871, and an NIH training grant T32 GM008759. We acknowledge the BioFrontiers Computing Core at the University of Colorado Boulder for providing highperformance computing resources (NIH 1S10OD012300) supported by BioFrontiers’ IT. In particular we thank Matt Hynes-Grace, Jon DeMasi, and Ethan Kern for assistance in the development of the TFEA website. Finally, we also thank the BioFrontiers Institute Next-Gen Sequencing Core and the Biochemistry Shared Cell Culture Facility for their invaluable contributions to this study.

## Notes

#### Summary of Updates

Supplemental file and acknowledgements section added. Link to TFEA website fixed

