## Supplementary material for "Transcription factor enrichment analysis (TFEA): Quantifying the activity of hundreds of transcription factors from a single experiment": All Supplemental Material

### 1 Accession Tables

#### 1.1 Figure 3 Accession Table

| Target | Treatment | Data Type | Cell Type | Accession |
|---|---|---|---|---|
| Nascent RNA | DMSO 1hr | GRO-seq | HCT116 | SRR1105736 |
| Nascent RNA | DMSO 1hr | GRO-seq | HCT116 | SRR1105737 |
| Nascent RNA | Nutlin 1hr | GRO-seq | HCT116 | SRR1105738 |
| Nascent RNA | Nutlin 1hr | GRO-seq | HCT116 | SRR1105739 |

Supplemental Table 1: Accession numbers used in Figure 3, data from [1].

#### 1.2 Figure 5 Accession Table

| Target | Treatment | Data Type | Cell Type | Accession |
|---|---|---|---|---|
| Capped RNA | LPS 0hr | CAGE | Macrophage donor2 | 12796-136F6 |
| Capped RNA | LPS 0.25hr | CAGE | Macrophage donor2 | 12797-136F7 |
| Capped RNA | LPS 0.5hr | CAGE | Macrophage donor2 | 12798-136F8 |
| Capped RNA | LPS 0.75hr | CAGE | Macrophage donor2 | 12799-136F9 |
| Capped RNA | LPS 1hr | CAGE | Macrophage donor2 | 12800-136G1 |
| Capped RNA | LPS 1.3hr | CAGE | Macrophage donor2 | 12801-136G2 |
| Capped RNA | LPS 2hr | CAGE | Macrophage donor2 | 12803-136G4 |
| Capped RNA | LPS 2.5hr | CAGE | Macrophage donor2 | 12804-136G5 |
| Capped RNA | LPS 3hr | CAGE | Macrophage donor2 | 12805-136G6 |
| Capped RNA | LPS 3.5hr | CAGE | Macrophage donor2 | 12806-136G7 |
| Capped RNA | LPS 4hr | CAGE | Macrophage donor2 | 12807-136G8 |
| Capped RNA | LPS 5hr | CAGE | Macrophage donor2 | 12808-136G9 |
| Capped RNA | LPS 8hr | CAGE | Macrophage donor2 | 12811-136H3 |
| Capped RNA | LPS 10hr | CAGE | Macrophage donor2 | 12812-136H4 |
| Capped RNA | LPS 12hr | CAGE | Macrophage donor2 | 12813-136H5 |
| Capped RNA | LPS 14hr | CAGE | Macrophage donor2 | 12814-136H6 |
| Capped RNA | LPS 16hr | CAGE | Macrophage donor2 | 12815-136H7 |
| Capped RNA | LPS 18hr | CAGE | Macrophage donor2 | 12816-136H8 |
| Capped RNA | LPS 20hr | CAGE | Macrophage donor2 | 12817-136H9 |
| Capped RNA | LPS 22hr | CAGE | Macrophage donor2 | 12818-136I1 |
| Capped RNA | LPS 24hr | CAGE | Macrophage donor2 | 12819-136I2 |
| Capped RNA | LPS 36hr | CAGE | Macrophage donor2 | 12820-136I3 |
| Capped RNA | LPS 48hr | CAGE | Macrophage donor2 | 12821-136I4 |

Supplemental Table 2: Project numbers used in Figure 5, data from [6, 4].

#### 1.3 Figure 6 Accession Table

| Target | Treatment | Data Type | Cell Type | Accession |
|---|---|---|---|---|
| ATAC | Dex 0hr | ATAC-seq | A549 | ENCSR220ASC |
| ATAC | Dex 1hr | ATAC-seq | A549 | ENCSR139OYS |
| ATAC | Dex 4hr | ATAC-seq | A549 | ENCSR288YMH |
| ATAC | Dex 8hr | ATAC-seq | A549 | ENCSR074AHH |

| | | | | |
|---|---|---|---|---|
| ATAC | Dex 12hr | ATAC-seq | A549 | ENCSR265ZXX |
| DNase | Dex 0hr | DNase-seq | A549 | ENCSR136DNA |
| DNase | Dex 30min | DNase-seq | A549 | ENCSR406EMB |
| DNase | Dex 1hr | DNase-seq | A549 | ENCSR384KCZ |
| DNase | Dex 2hr | DNase-seq | A549 | ENCSR837VHE |
| DNase | Dex 3hr | DNase-seq | A549 | ENCSR294XUZ |
| DNase | Dex 4hr | DNase-seq | A549 | ENCSR599WJC |
| DNase | Dex 5hr | DNase-seq | A549 | ENCSR565WPR |
| DNase | Dex 6hr | DNase-seq | A549 | ENCSR077EYC |
| DNase | Dex 7hr | DNase-seq | A549 | ENCSR347CEH |
| DNase | Dex 8hr | DNase-seq | A549 | ENCSR660OQE |
| DNase | Dex 10hr | DNase-seq | A549 | ENCSR128IVG |
| DNase | Dex 12hr | DNase-seq | A549 | ENCSR523FJT |
| EP300 | Dex 0hr | ChIP-seq | A549 | ENCSR886OEO |
| EP300 | Dex 5min | ChIP-seq | A549 | ENCSR602BTS |
| EP300 | Dex 10min | ChIP-seq | A549 | ENCSR174FJD |
| EP300 | Dex 15min | ChIP-seq | A549 | ENCSR788VKG |
| EP300 | Dex 20min | ChIP-seq | A549 | ENCSR167QIJ |
| EP300 | Dex 25min | ChIP-seq | A549 | ENCSR044IFH |
| EP300 | Dex 30min | ChIP-seq | A549 | ENCSR260WCE |
| EP300 | Dex 1hr | ChIP-seq | A549 | ENCSR358ELZ |
| EP300 | Dex 2hr | ChIP-seq | A549 | ENCSR770OTI |
| EP300 | Dex 3hr | ChIP-seq | A549 | ENCSR047EVQ |
| EP300 | Dex 4hr | ChIP-seq | A549 | ENCSR145YCX |
| EP300 | Dex 5hr | ChIP-seq | A549 | ENCSR610RKF |
| EP300 | Dex 6hr | ChIP-seq | A549 | ENCSR841ASB |
| EP300 | Dex 7hr | ChIP-seq | A549 | ENCSR467VXG |
| EP300 | Dex 8hr | ChIP-seq | A549 | ENCSR561ZRE |
| EP300 | Dex 10hr | ChIP-seq | A549 | ENCSR792VMN |
| EP300 | Dex 12hr | ChIP-seq | A549 | ENCSR124VXG |
| H3K27ac | Dex 0hr | ChIP-seq | A549 | ENCSR778NQS |
| H3K27ac | Dex 5min | ChIP-seq | A549 | ENCSR734FLK |
| H3K27ac | Dex 10min | ChIP-seq | A549 | ENCSR027BPE |
| H3K27ac | Dex 15min | ChIP-seq | A549 | ENCSR325VCV |
| H3K27ac | Dex 20min | ChIP-seq | A549 | ENCSR864KVZ |
| H3K27ac | Dex 25min | ChIP-seq | A549 | ENCSR480OHP |

| | | | | |
|---|---|---|---|---|
| H3K27ac | Dex 30min | ChIP-seq | A549 | ENCSR102XUM |
| H3K27ac | Dex 1hr | ChIP-seq | A549 | ENCSR242TBH |
| H3K27ac | Dex 2hr | ChIP-seq | A549 | ENCSR614NPG |
| H3K27ac | Dex 3hr | ChIP-seq | A549 | ENCSR350EFV |
| H3K27ac | Dex 4hr | ChIP-seq | A549 | ENCSR543ZVZ |
| H3K27ac | Dex 5hr | ChIP-seq | A549 | ENCSR716XDB |
| H3K27ac | Dex 6hr | ChIP-seq | A549 | ENCSR340NAL |
| H3K27ac | Dex 7hr | ChIP-seq | A549 | ENCSR569IBY |
| H3K27ac | Dex 8hr | ChIP-seq | A549 | ENCSR250EHC |
| H3K27ac | Dex 10hr | ChIP-seq | A549 | ENCSR180YHA |
| H3K27ac | Dex 12hr | ChIP-seq | A549 | ENCSR435JKM |
| H3K4me1 | Dex 0hr | ChIP-seq | A549 | ENCSR636PIN |
| H3K4me1 | Dex 30min | ChIP-seq | A549 | ENCSR593RGY |
| H3K4me1 | Dex 1hr | ChIP-seq | A549 | ENCSR537FVU |
| H3K4me1 | Dex 2hr | ChIP-seq | A549 | ENCSR726MAP |
| H3K4me1 | Dex 3hr | ChIP-seq | A549 | ENCSR171ZJG |
| H3K4me1 | Dex 4hr | ChIP-seq | A549 | ENCSR949IDI |
| H3K4me1 | Dex 5hr | ChIP-seq | A549 | ENCSR225AOO |
| H3K4me1 | Dex 6hr | ChIP-seq | A549 | ENCSR868MLT |
| H3K4me1 | Dex 7hr | ChIP-seq | A549 | ENCSR462JVS |
| H3K4me1 | Dex 8hr | ChIP-seq | A549 | ENCSR404OLV |
| H3K4me1 | Dex 10hr | ChIP-seq | A549 | ENCSR954HUB |
| H3K4me1 | Dex 12hr | ChIP-seq | A549 | ENCSR529YKU |
| H3K4me2 | Dex 0hr | ChIP-seq | A549 | ENCSR410BCN |
| H3K4me2 | Dex 30min | ChIP-seq | A549 | ENCSR215DID |
| H3K4me2 | Dex 1hr | ChIP-seq | A549 | ENCSR692JHM |
| H3K4me2 | Dex 2hr | ChIP-seq | A549 | ENCSR124YCC |
| H3K4me2 | Dex 3hr | ChIP-seq | A549 | ENCSR918VQU |
| H3K4me2 | Dex 4hr | ChIP-seq | A549 | ENCSR834LCU |
| H3K4me2 | Dex 5hr | ChIP-seq | A549 | ENCSR555EAA |
| H3K4me2 | Dex 6hr | ChIP-seq | A549 | ENCSR905REY |
| H3K4me2 | Dex 7hr | ChIP-seq | A549 | ENCSR016PSC |
| H3K4me2 | Dex 8hr | ChIP-seq | A549 | ENCSR774KCU |
| H3K4me2 | Dex 10hr | ChIP-seq | A549 | ENCSR766NHB |
| H3K4me2 | Dex 12hr | ChIP-seq | A549 | ENCSR428DFL |
| H3K4me3 | Dex 0hr | ChIP-seq | A549 | ENCSR203XPU |

| | | | | |
|---|---|---|---|---|
| H3K4me3 | Dex 30min | ChIP-seq | A549 | ENCSR677QYM |
| H3K4me3 | Dex 1hr | ChIP-seq | A549 | ENCSR928FDN |
| H3K4me3 | Dex 2hr | ChIP-seq | A549 | ENCSR252FZA |
| H3K4me3 | Dex 3hr | ChIP-seq | A549 | ENCSR483JJT |
| H3K4me3 | Dex 4hr | ChIP-seq | A549 | ENCSR901AAW |
| H3K4me3 | Dex 5hr | ChIP-seq | A549 | ENCSR524UOX |
| H3K4me3 | Dex 6hr | ChIP-seq | A549 | ENCSR646OPC |
| H3K4me3 | Dex 7hr | ChIP-seq | A549 | ENCSR285FZP |
| H3K4me3 | Dex 8hr | ChIP-seq | A549 | ENCSR618MUP |
| H3K4me3 | Dex 10hr | ChIP-seq | A549 | ENCSR139DGM |
| H3K4me3 | Dex 12hr | ChIP-seq | A549 | ENCSR944WVU |
| H3K9me3 | Dex 0hr | ChIP-seq | A549 | ENCSR775TAI |
| H3K9me3 | Dex 30min | ChIP-seq | A549 | ENCSR109KEL |
| H3K9me3 | Dex 1hr | ChIP-seq | A549 | ENCSR954SQX |
| H3K9me3 | Dex 2hr | ChIP-seq | A549 | ENCSR936UEX |
| H3K9me3 | Dex 3hr | ChIP-seq | A549 | ENCSR354ERB |
| H3K9me3 | Dex 4hr | ChIP-seq | A549 | ENCSR037DAC |
| H3K9me3 | Dex 5hr | ChIP-seq | A549 | ENCSR299MNA |
| H3K9me3 | Dex 6hr | ChIP-seq | A549 | ENCSR032SCO |
| H3K9me3 | Dex 7hr | ChIP-seq | A549 | ENCSR873VJA |
| H3K9me3 | Dex 8hr | ChIP-seq | A549 | ENCSR791BNU |
| H3K9me3 | Dex 10hr | ChIP-seq | A549 | ENCSR013RTF |
| H3K9me3 | Dex 12hr | ChIP-seq | A549 | ENCSR451MJX |

Supplemental Table 3: Accession numbers used in Figure 6, data from [5, 8].

**DMSO vs. Nutlin TFEA Results**

a

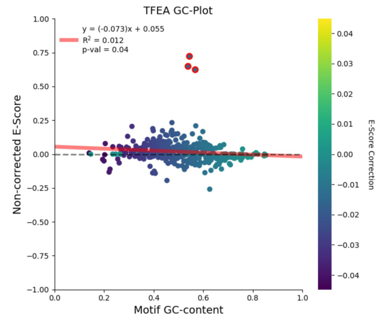


b

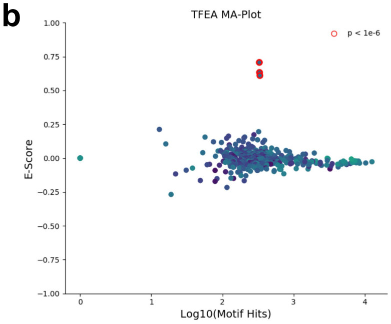


c

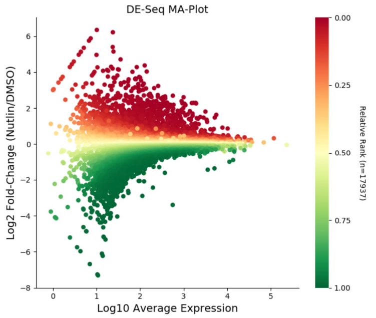


d

**p-adj < 1e-6**

User Inputs | Text Results

MD Results | MD MA-Plot | MD VolcanoPlot

MDD Results | MDD MA-Plot | MDD VolcanoPlot

| Module | Value | Time (hh:mm:ss) |
|---|---|---|
| COMBINE | mumerge | 0:10:48 |
| RANK | deseq | 1:53:05 |
| SCANNER | fimo | 0:09:54 |
| ENRICHMENT | auc | 0:04:42 |
| OUTPUT | html | 0:00:02 |
| **Total** | | **2:18:33** |

e

**Positive E-Score**

| #TF | E-Score | Corrected E-Score | Events | GC | FPKM | P-adj | Corrected P-adj |
|---|---|---|---|---|---|---|---|
| P53_HUMAN.H11MO.0.A | 0.722 | 0.707 | 331 | 0.545 | 9.07 | 1e-40 | 1e-39 |
| P73_HUMAN.H11MO.0.A | 0.648 | 0.633 | 333 | 0.539 | 2.19 | 1e-33 | 1e-31 |
| P63_HUMAN.H11MO.0.A | 0.624 | 0.61 | 337 | 0.568 | 0.0492 | 1e-31 | 1e-29 |
| VDR_HUMAN.H11MO.0.A | 0.209 | 0.195 | 324 | 0.57 | 2.42 | 1e-2 | 1e-1 |
| FOXC1_HUMAN.H11MO.0.C | 0.205 | 0.173 | 278 | 0.313 | 30.4 | 1e-2 | 1e-1 |
| BACH2_HUMAN.H11MO.0.A | 0.151 | 0.132 | 393 | 0.497 | 0.519 | 0.0752 | 0.207 |
| FOSL1_HUMAN.H11MO.0.A | 0.163 | 0.143 | 267 | 0.471 | 123 | 0.136 | 0.337 |
| SMAD4_HUMAN.H11MO.0.B | 0.15 | 0.135 | 313 | 0.557 | 8.43 | 0.186 | 0.352 |
| JUND_HUMAN.H11MO.0.A | 0.176 | 0.156 | 206 | 0.474 | 183 | 0.277 | 0.55 |
| JUNB_HUMAN.H11MO.0.A | 0.138 | 0.116 | 372 | 0.454 | 465 | 0.202 | 0.555 |
| MAFK_HUMAN.H11MO.0.A | 0.173 | 0.149 | 201 | 0.436 | 11.8 | 0.351 | 0.723 |
| CUX1_HUMAN.H11MO.0.C | 0.233 | 0.212 | 13 | 0.474 | 4.5 | 1 | 1 |
| CEBPG_HUMAN.H11MO.0.B | 0.188 | 0.162 | 122 | 0.397 | 13.7 | 0.669 | 1 |
| MTF1_HUMAN.H11MO.0.C | 0.167 | 0.155 | 57 | 0.599 | 5.45 | 1 | 1 |
| ATF4_HUMAN.H11MO.0.A | 0.173 | 0.147 | 115 | 0.399 | 517 | 1 | 1 |

⋮

**Negative E-Score**

| #TF | E-Score | Corrected E-Score | Events | GC | FPKM | P-adj | Corrected P-adj |
|---|---|---|---|---|---|---|---|
| ZN121_HUMAN.H11MO.0.C | -0.0234 | -0.0359 | 1.04e+04 | 0.583 | 27.4 | 1e-1 | 1e-5 |
| FOXJ3_HUMAN.H11MO.0.A | -0.0391 | -0.0799 | 3.27e+03 | 0.195 | 4.8 | 0.286 | 1e-4 |
| ZN770_HUMAN.H11MO.0.C | -0.0205 | -0.027 | 1.27e+04 | 0.664 | 36.5 | 1e-2 | 1e-4 |
| THA_HUMAN.H11MO.0.C | -0.0492 | -0.0653 | 4.26e+03 | 0.534 | 1.75 | 1e-2 | 1e-4 |
| PAX5_HUMAN.H11MO.0.A | -0.0283 | -0.0374 | 8.51e+03 | 0.629 | 0.179 | 1e-1 | 1e-3 |
| PRDM6_HUMAN.H11MO.0.C | -0.0162 | -0.0475 | 5.59e+03 | 0.325 | 0.22 | 1 | 1e-3 |
| ZFP28_HUMAN.H11MO.0.C | -0.029 | -0.0588 | 3.36e+03 | 0.346 | 0.0293 | 0.841 | 1e-2 |
| IRF3_HUMAN.H11MO.0.B | -0.0158 | -0.0401 | 6.3e+03 | 0.421 | 11.4 | 1 | 1e-2 |
| VEZF1_HUMAN.H11MO.0.C | -0.0257 | -0.0343 | 7.64e+03 | 0.636 | 26.9 | 0.115 | 1e-2 |
| SP4_HUMAN.H11MO.0.A | -0.0397 | -0.0457 | 4.74e+03 | 0.672 | 1.95 | 1e-1 | 1e-2 |
| ZBT17_HUMAN.H11MO.0.A | -0.0331 | -0.0423 | 5.46e+03 | 0.627 | 3.62 | 0.121 | 1e-1 |
| ZSC22_HUMAN.H11MO.0.C | -0.0741 | -0.0806 | 1.6e+03 | 0.664 | 7.11 | 0.0566 | 1e-1 |
| KLF15_HUMAN.H11MO.0.A | -0.0298 | -0.032 | 6.79e+03 | 0.724 | 1.31 | 0.0833 | 1e-1 |
| WT1_HUMAN.H11MO.0.C | -0.0304 | -0.0323 | 6.68e+03 | 0.728 | 0.0377 | 0.0799 | 1e-1 |
| ZN281_HUMAN.H11MO.0.A | -0.0445 | -0.0446 | 4.19e+03 | 0.752 | 24.3 | 1e-1 | 1e-1 |

⋮

Supplemental Figure 1: **An example of TFEA HTML results page.** (a) Pre-GC correction showing the E-Score of each motif (y-axis) as a function of GC-content (x-axis). Red line: linear regression fit; dots colored by the amount to correct. (b) A scatter plot (colored as in a), similar to an MA-plot, showing the GC-corrected E-Scores (y-axis) vs number of motif hits within regions (<1.5kb; x-axis) for each motif analyzed. (c) An MA-plot of the ROIs generated from DE-Seq2. (d) A table listing the inputs, text results, MD-Score and MDD-Score results (as clickable links), as well as the time taken to complete each step of the TFEA process. (e) A list of motifs that exhibit positive or negative enrichment ordered by adjusted p-value. Significant motifs appear as red and have clickable links to individual results pages with more detailed information. Data is HCT116 dataset, as used in Figure 1a[1].

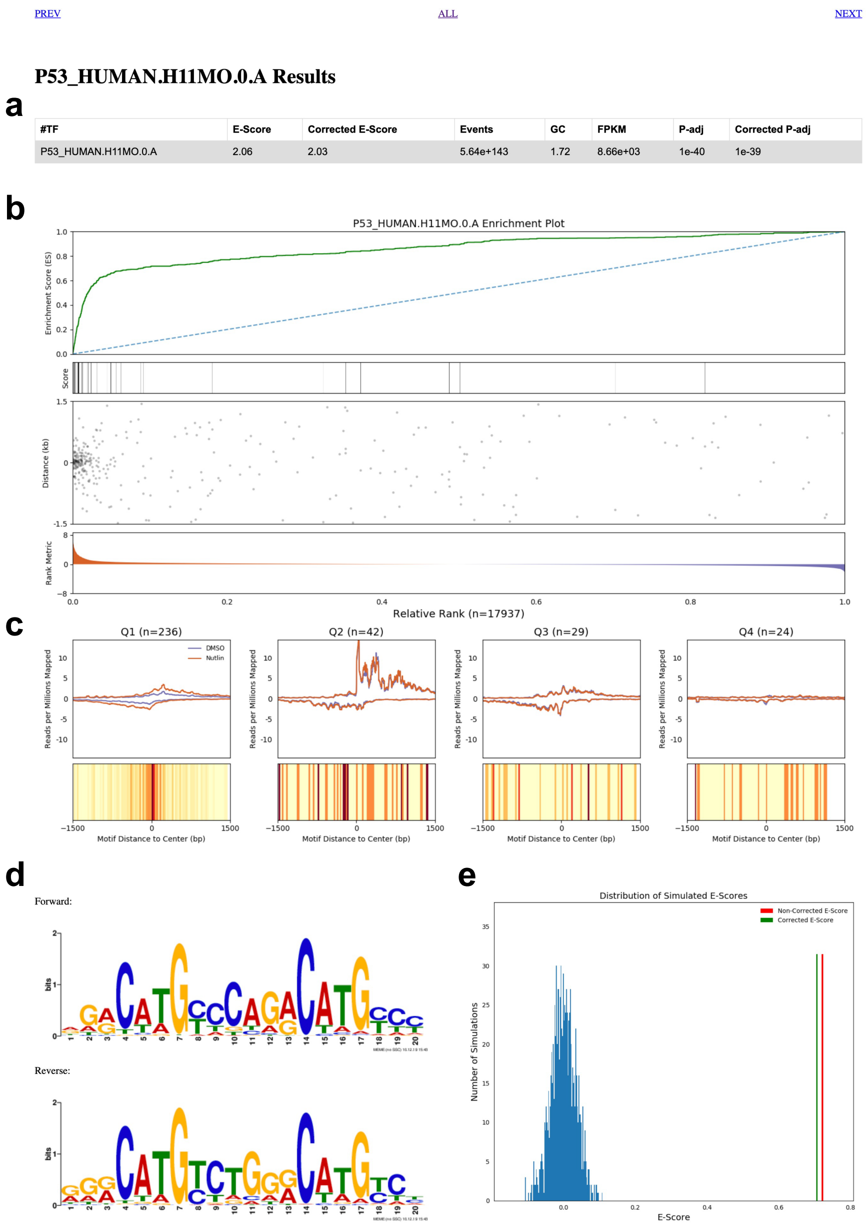


Supplemental Figure 2: **An example of a TFEA individual motif results page.** This page is reached by clicking on the corresponding motif in Figure 1e. (a) Summary statistics for the motif of interest, in this case p53 from HOCOMOCO v11. (b) Enrichment plot showing (from top to bottom) the running sum statistic (green line), the individual scores of each ROI (as heatmap), scatter plot of motif hits within ROIs relative to the reference point (labeled 0), and the ranking of ROIs based on differential transcription (red: positive; blue: negative). (c) For each quartile, summarize motif containing ROI within the quartile via Top: Meta plot of read coverage over ROIs. Bottom: Motif displacement distribution (as heatmap) summarizing the motif positions relative to the reference point. (Yellow is background to Red at max instances). (d) Forward and reverse complement position specific scoring matrix of the motif analyzed. (e) Histogram of E-Scores from randomly shuffling the rank order of ROIs (blue) with true non-corrected E-score (red) and GC-corrected E-score (green).

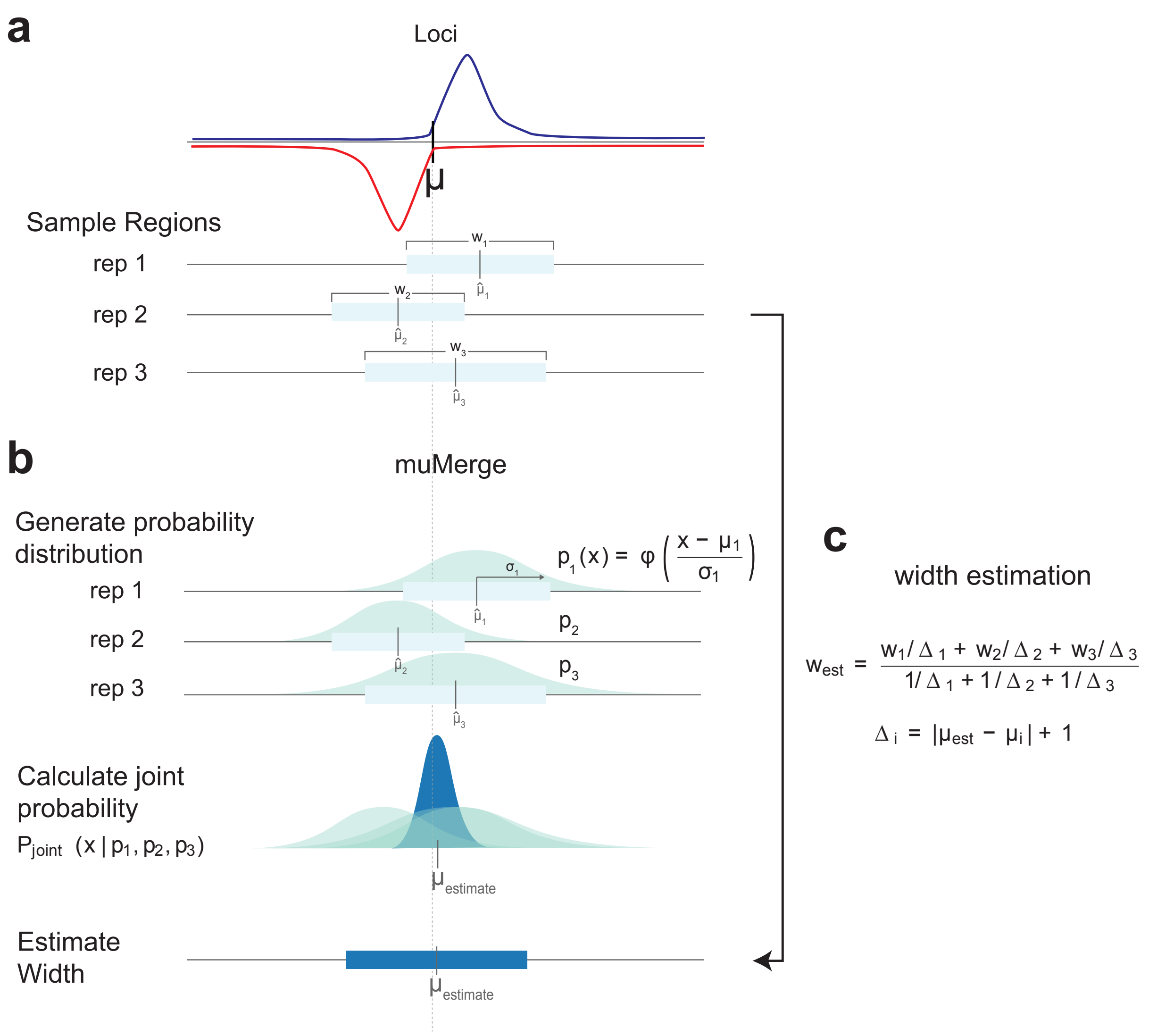


Supplemental Figure 3: **Diagrammatic description of the *muMerge* method** (a) The goal of *muMerge* is to combine multiple sample regions (light blue boxes) which originate from different replicates and/or conditions, but are measurements of the same underlying loci $\mu$, into a consensus set of ROIs. Red and blue lines are hypothetical data. (b) *muMerge* assumes that each sample region is an estimate on the location of a genomic loci of interest and models this probability ($p_i$) as a normal distribution (light blue) with $\mu_i$ equal to the center of each sample region. Subsequently, a joint probability ($p_{joint}$, dark blue) is calculated from the samples, and the estimate for the consensus position ($\mu_{estimate}$) is the maxima of this joint distribution. (c) Finally, to calculate the best estimate for the width of the ROI, a weighted average of the sample region widths is calculated. It is assumed that the sample regions closest to the consensus position are the most accurate representation of the underlying loci, so the weighted average of the widths is calculated such that more weight is given to the sample regions closer to $\mu_{estimate}$.

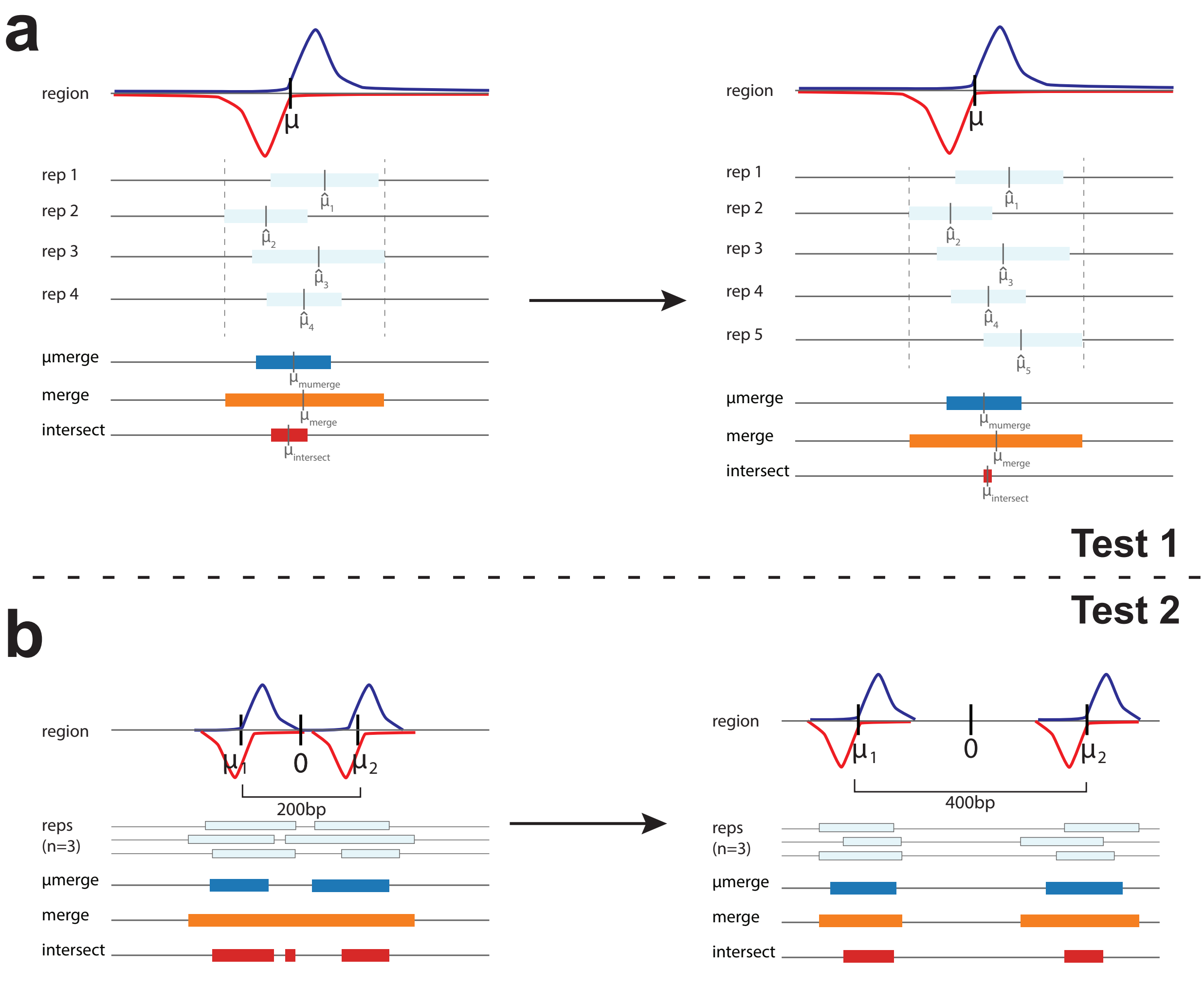


Supplemental Figure 4: **Tests performed to compare the performance of *muMerge* with bedtools merge and bedtools intersect.** (a) The first test involves sampling regions from a single theoretical loci with increasing replicates (light blue). *muMerge* retains correct length and *mu* position, bedtools merge tends to increase ROI length and bedtools intersect tends to decrease ROI length as more replicates are included. (b) A second test to determine performance when sampling from two theoretical loci as a function of inter-loci spacing. For closely spaced loci, *muMerge* correctly separates the two loci whereas bedtools merge is more likely to generate a single ROI, and bedtools intersect is more likely to generate multiple separate ROI (in this example, three). For both tests, top cartoon depicts sequencing data histograms on two strands (blue: positive strand; red (negative strand). Regions inferred from individual replicates in light blue. ROI ascertained by *muMerge* (dark blue), *bedtools merge* (orange) and *bedtools intersect* (red) shown for comparison.

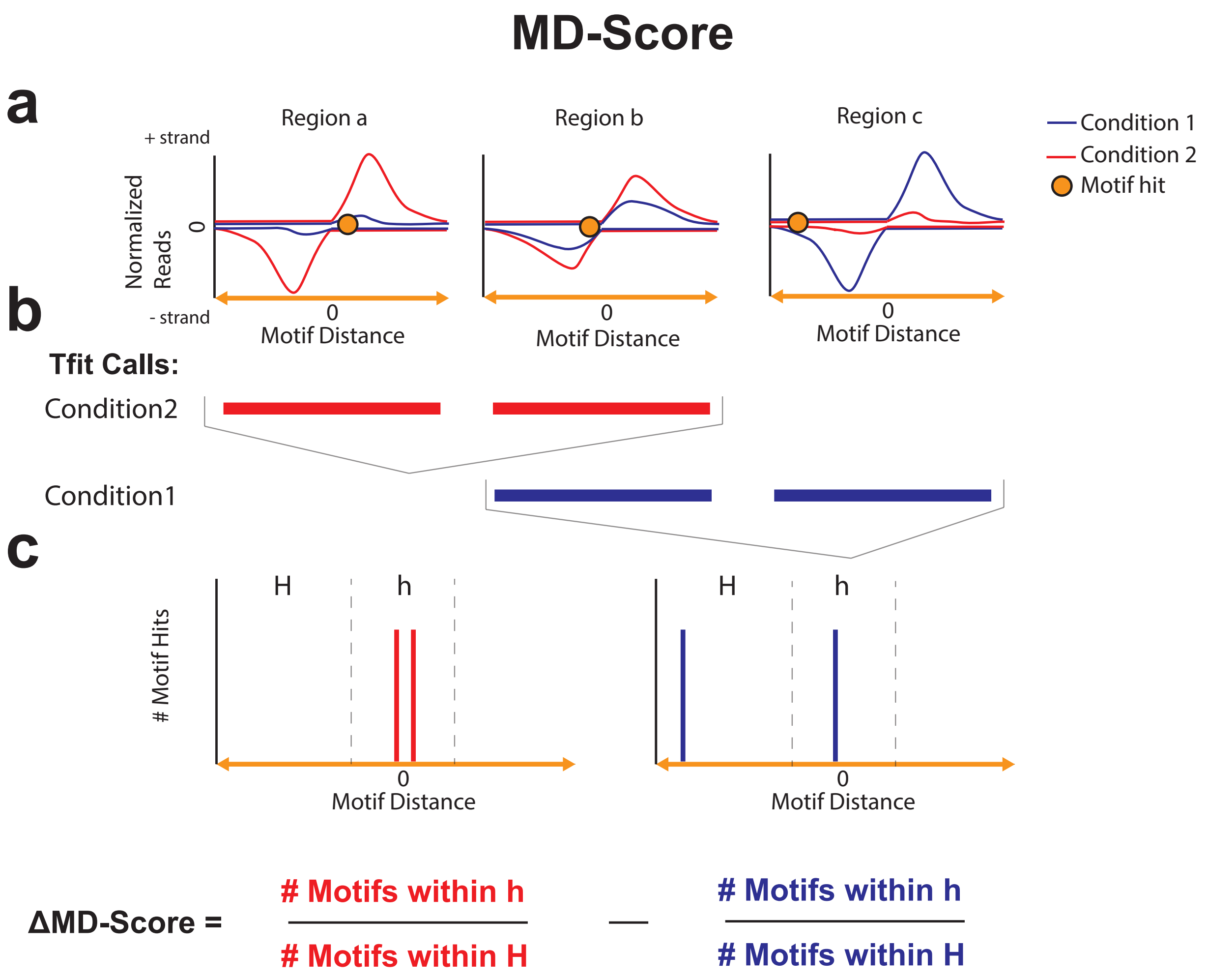


Supplemental Figure 5: **Cartoon diagram description of MD-Score method.** (a) Cartoon depicting typical histograms of nascent transcription data for three example regions. Orange dot is motif location. (b) Tfit called sites of RNA polymerase initiation in each dataset (red, blue) as called by Tfit[3]. These regions are the inputs to the MD-score approach[2]. (c) Motif displacement distribution histograms plot position of motif (vertical bars) relative to reference point (labeled 0) for both conditions (red and blue). The MD-Score is the fraction of motif instances within the inner window (h) divided by the total motif hits in the larger window (H; note H encompasses h). MD-Scores are calculated independently in each of the two conditions to obtain the difference.

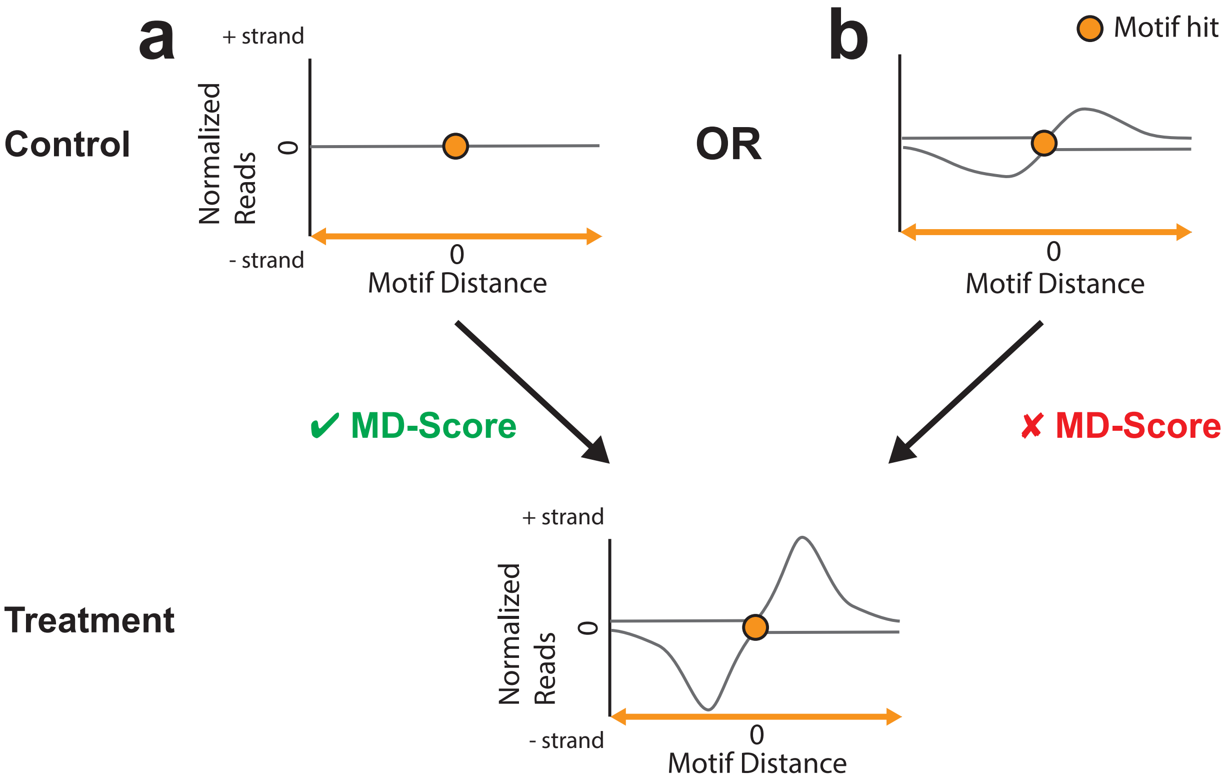


Supplemental Figure 6: **The MD-Score approach only detects gain or loss of transcribed regions.** A given locus in the treatment can arise from either (a) a region of no signal in the control; or (b) increase in signal at a pre-existing region within the control sample. Importantly, the first case increases the Δ MD-Score whereas the second does not alter the Δ MD-Score.

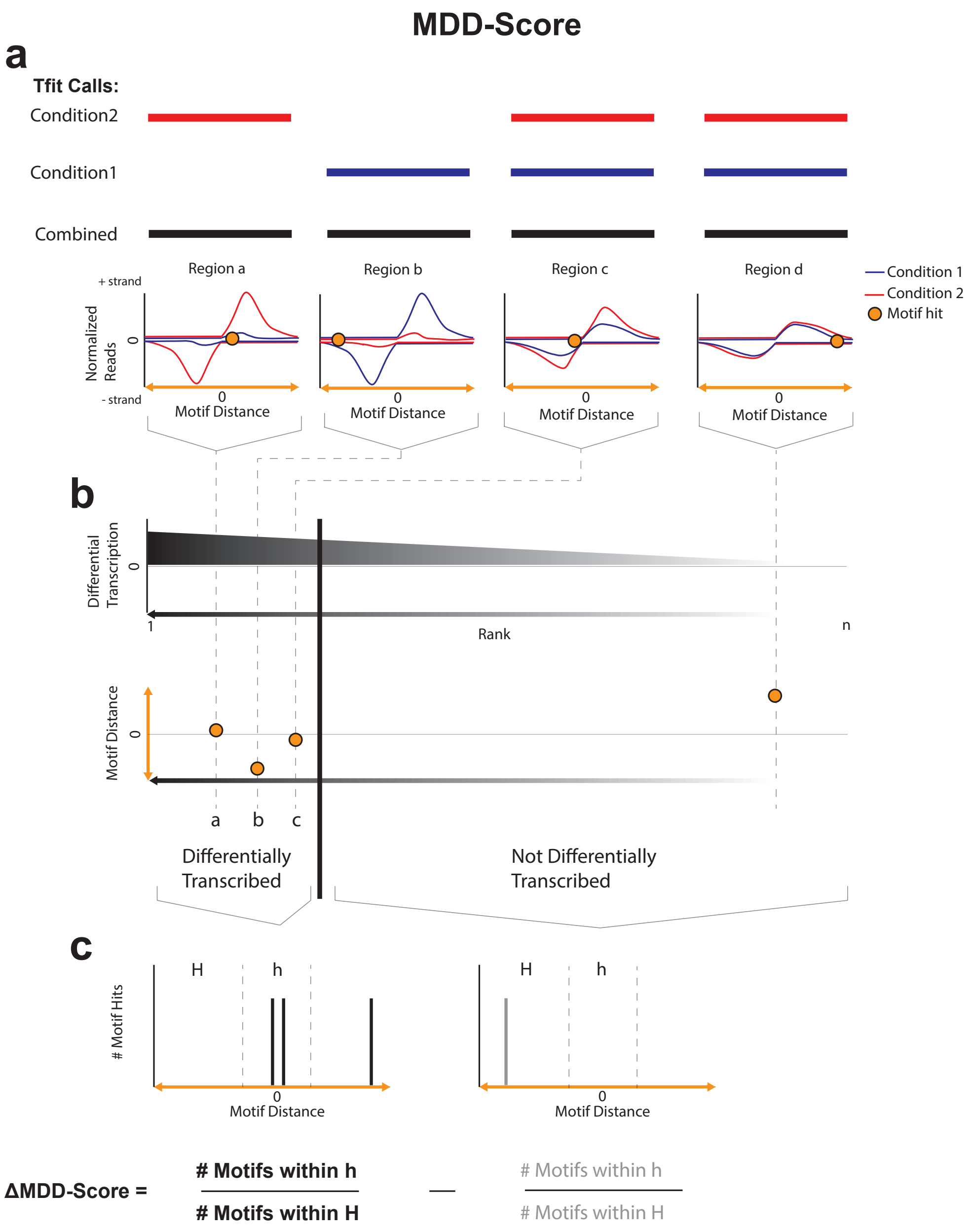


Supplemental Figure 7: **Cartoon diagram depicting the MDD-Score method.** The differential MD-Score method (referred to as MDD-Score)[9, 7] begins with (a) a collection of regions called in one or more conditions (red and blue). (b) Regions are ranked by DESeq or DESeq2 p-value (depending on replicate number) and a cutoff segregates identifies the differentially transcribed subset. (c) The differential MD-Score is calculated similarly to the MD-Score but between the differentially transcribed set (black) and the not differentially transcribed (in grey).

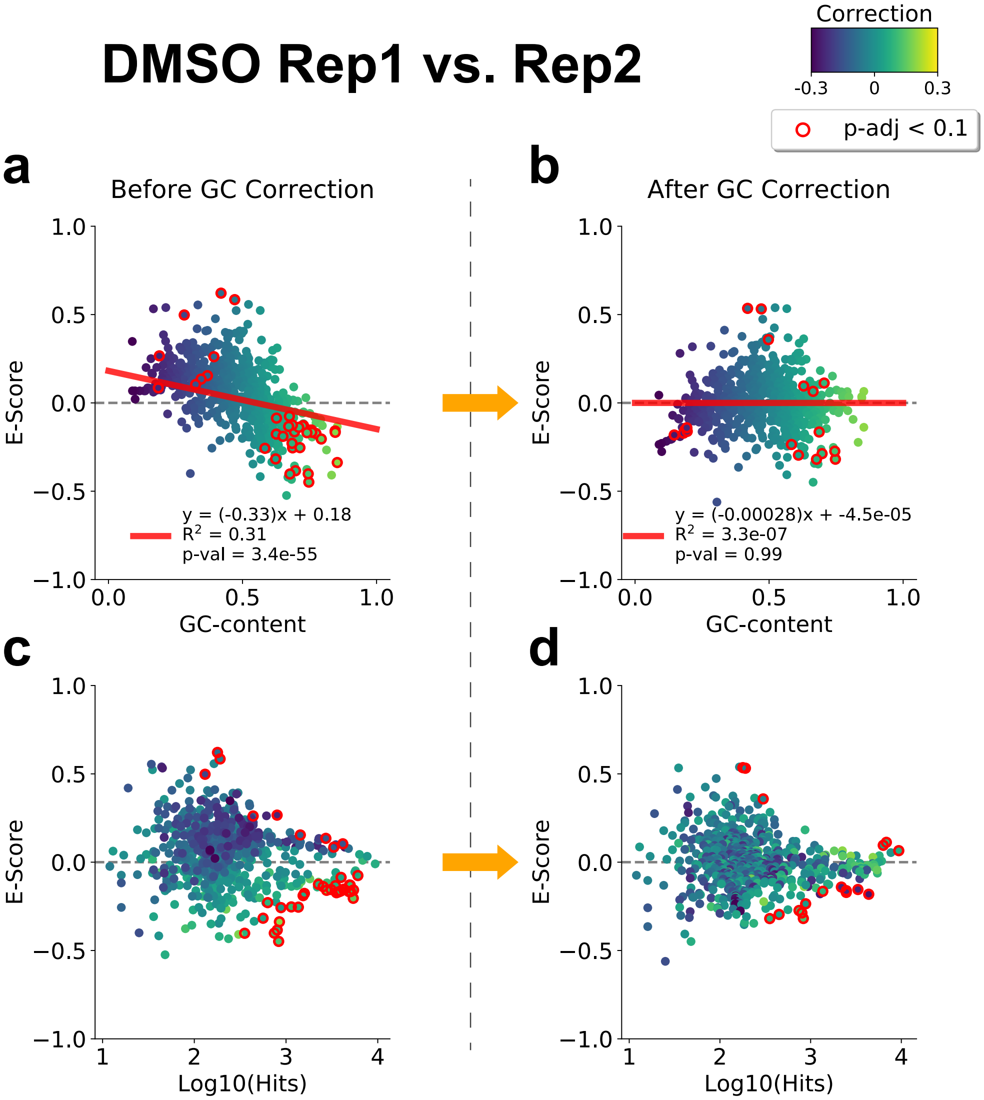


Supplemental Figure 8: **E-Scores are adjusted based on the GC content bias using linear regression.** We observed that motif E-Scores often correlated with their GC-content. (a) Scatter plot of E-Score (y-axis) vs. GC-content (x-axis) of motifs, comparing replicate 1 vs. replicate 2 (DMSO condition) before GC-correction (red line: linear regression fit). (b) Scatter plot of E-Score (y-axis) vs. GC-content (x-axis) of motifs after GC correction (red line: linear regression fit). (c) MA-plot of E-Score (y-axis) vs. Log10 of number of motif hits within regions of interest (x-axis) before GC correction. (d) MA-like plot of E-Score (y-axis) vs. Log10 of number of motif hits within regions of interest (x-axis) after GC-correction. These MA-Plots show that the underlying distribution of E-Scores relative to number of motif hits does not significantly change after GC-correction. All panels are data in HCT116 DMSO condition (SRR1105736, SRR1105737 [1], dots are colored by the amount to be corrected due to GC-bias, red outline dots are p-adj $< 0.1$).

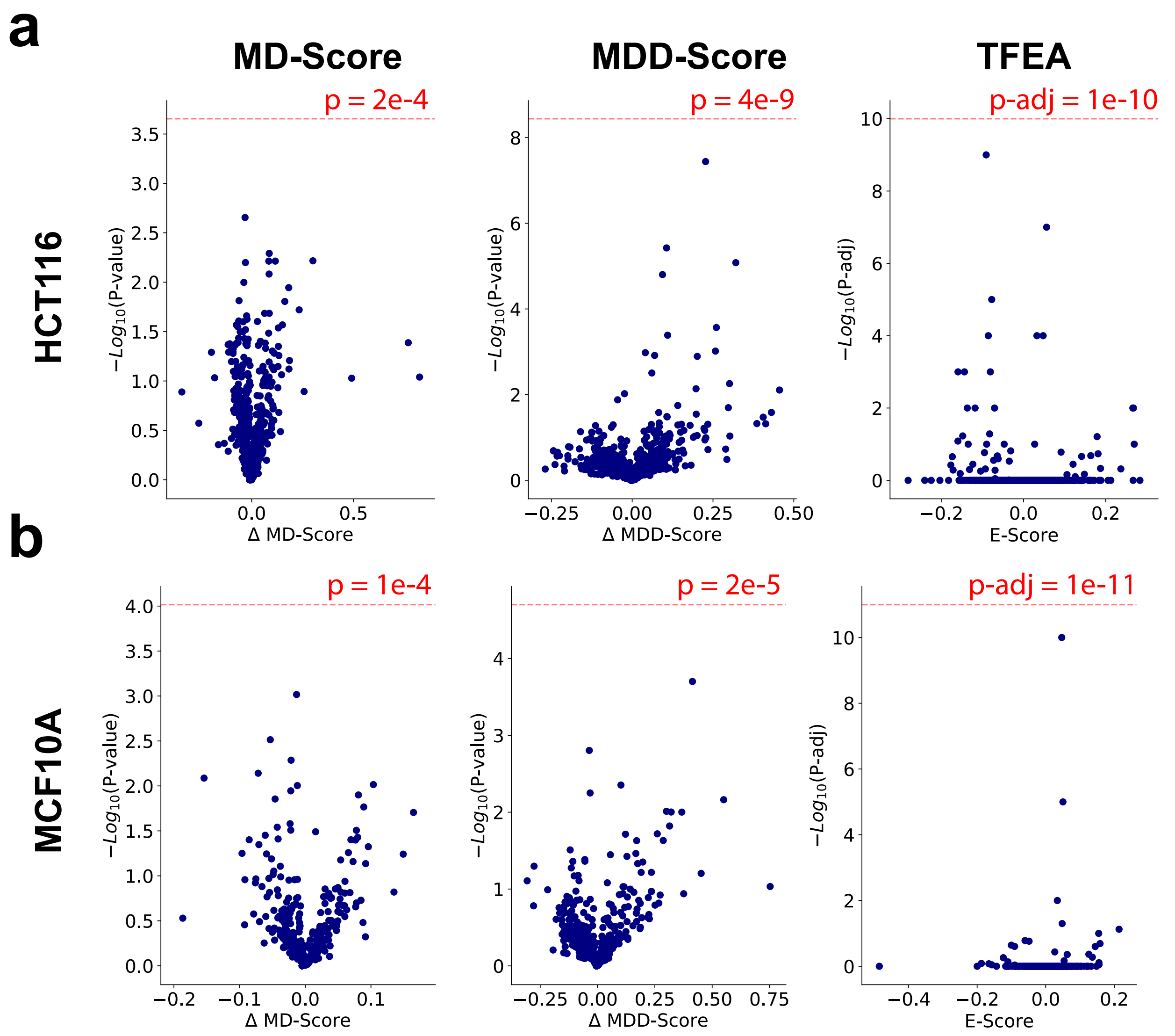


Supplemental Figure 9: **Choosing thresholds for MD-Score, MDD-Score, and TFEA.** To choose a threshold cutoff for the three methods used here, DMSO replicates were compared and the threshold at which no false positives are obtained was determined. To be conservative, an additional order of magnitude is added for stringency. We performed this for each method in either (a) HCT116 or (b) MCF10A cells.

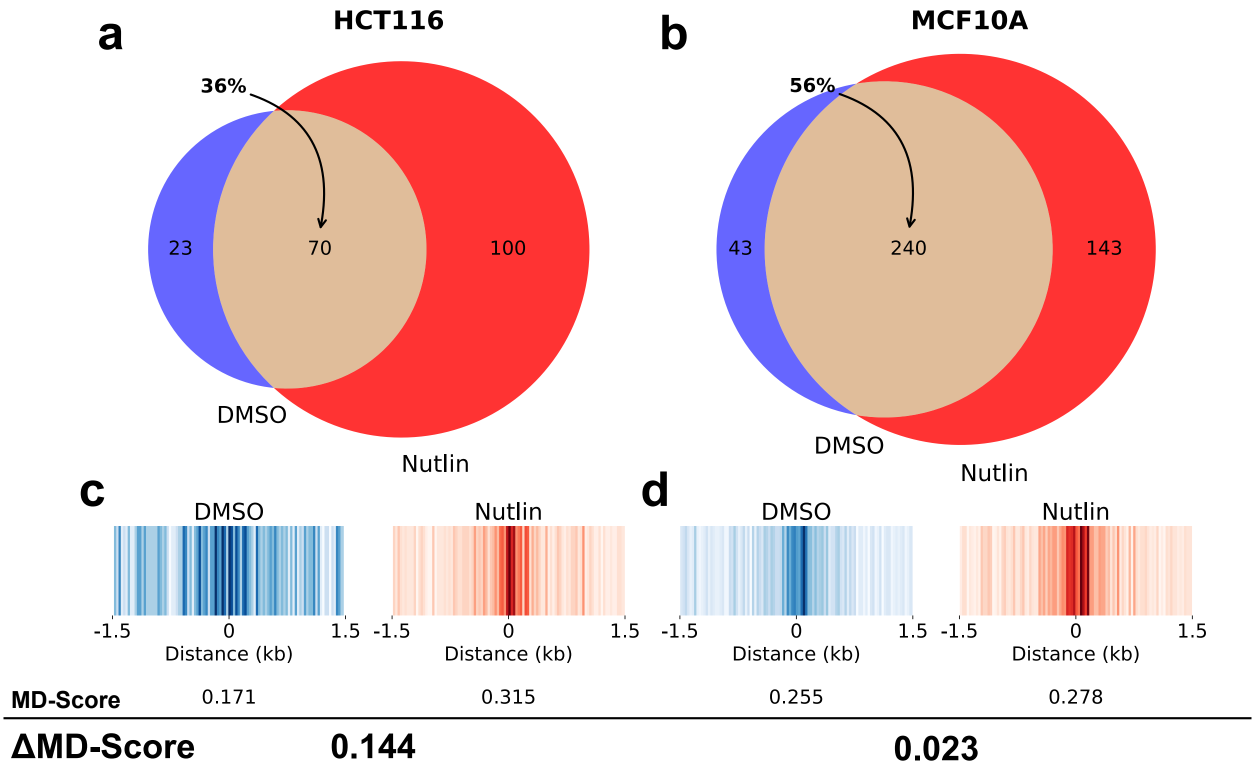


Supplemental Figure 10: **The MD-score approach fails to capture p53 after Nutlin treatment in MCF10A cells.** The response to Nutlin-3a visualized as Venn diagrams of (a) HCT116 and (b) MCF10a cells shows distinct p53 response, with a larger proportion (in MCF10A cells) of existing sites of RNA polymerase initiation that respond to Nutlin-3a. In both cases, only regions with p53 motif within 150 bps of the point of interest (midpoint of ROI) are shown. Motif displacement distributions of TP53 motif within 1.5 kb of ROI midpoints for (c) HCT116 or (d) MCF10A cells shows a higher co-localization of p53 in DMSO treated MCF10A cells. Bottom: MD-Score quantification for each condition followed by the observed ΔMD-Score for the Nutlin-3a response in each cell type.

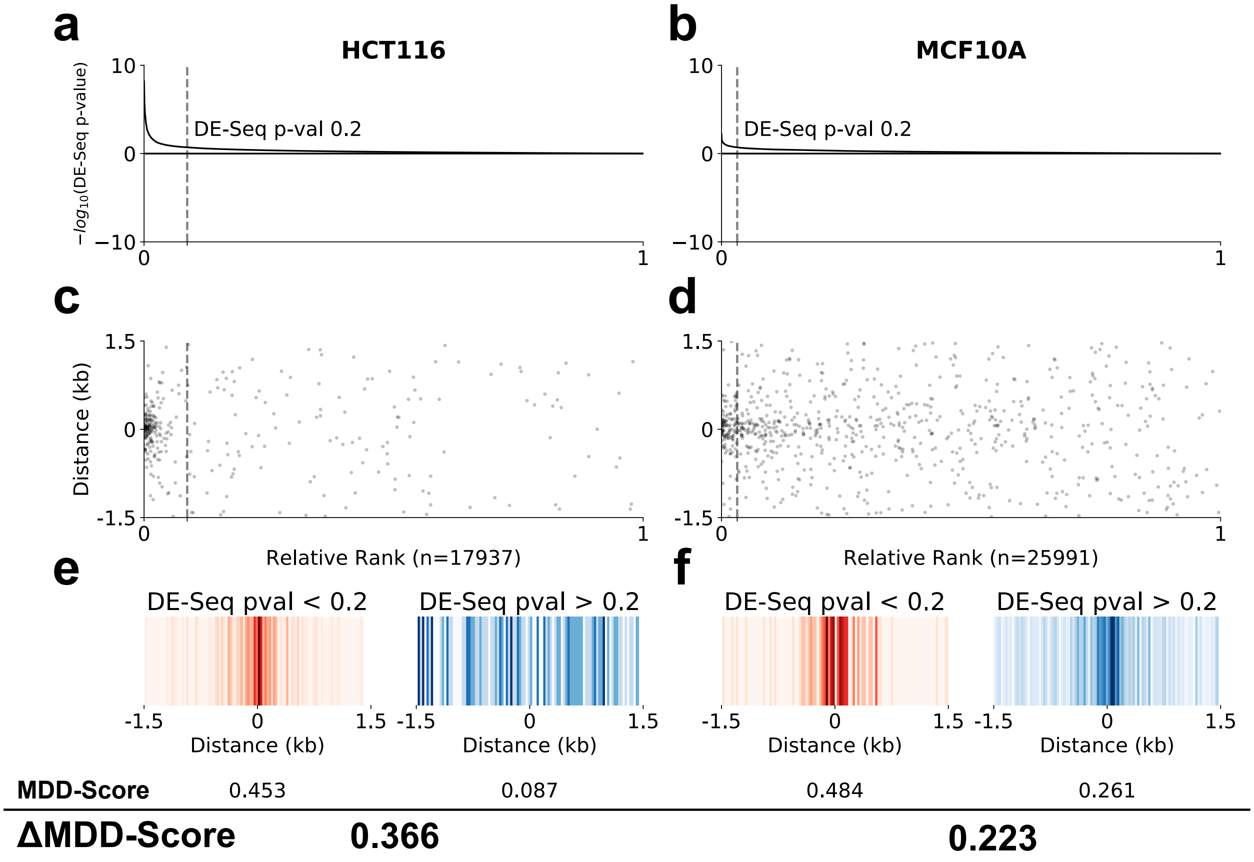


Supplemental Figure 11: **The MDD-Score method detects p53 following Nutlin treatment in both cell types.** The MDD-Score approach detects p53 response in both (a) HCT116 and (b) MCF10a cells. By default, a loose DESeq2 p-value of 0.2 is chosen to identify the set of differentially transcribed ROI. Scatterplots show instances of TP53 motif across ranked ROI for (c) HCT116 and (d) MCF10A cells. The presence of constitutive TP63 activity leads MCF10a cells to have a higher background signal around TP53 motifs. Motif displacement distribution heatmaps for (e) HCT116 and (f) MCF10A cells, further emphasize the increased background presence of the TP53 motifin MCF10A cells. Red is control (DMSO), blue is Nutlin treated. All panels are HCT116 data from SRR1105736, SRR1105737, SRR1105738, SRR1105739.

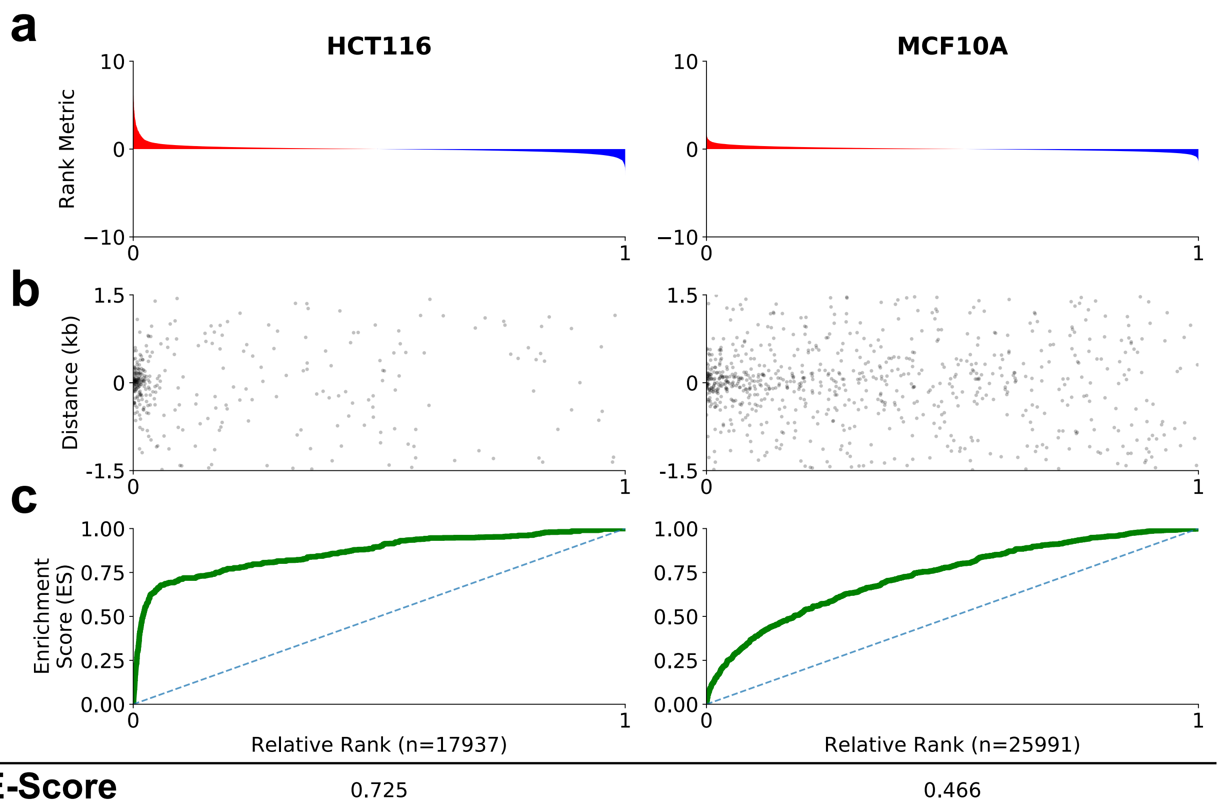


Supplemental Figure 12: **TFEA detects p53 in both HCT116 cells and MCF10A cells without the use of fixed thresholds.** (a) ROI are ranked by differential transcription. Red: increased transcription, blue: decreased. (b) Instances of the TP53 motif are detected within ranked ROIs. (c) TFEA measures motif enrichment as the E-Score, calculated as 2* AUC (ie. area under the curve) between the running sum of ROI scores (green line) and the uniform distribution (dashed blue line). HCT116 data from SRR1105736, SRR1105737, SRR1105738, SRR1105739.

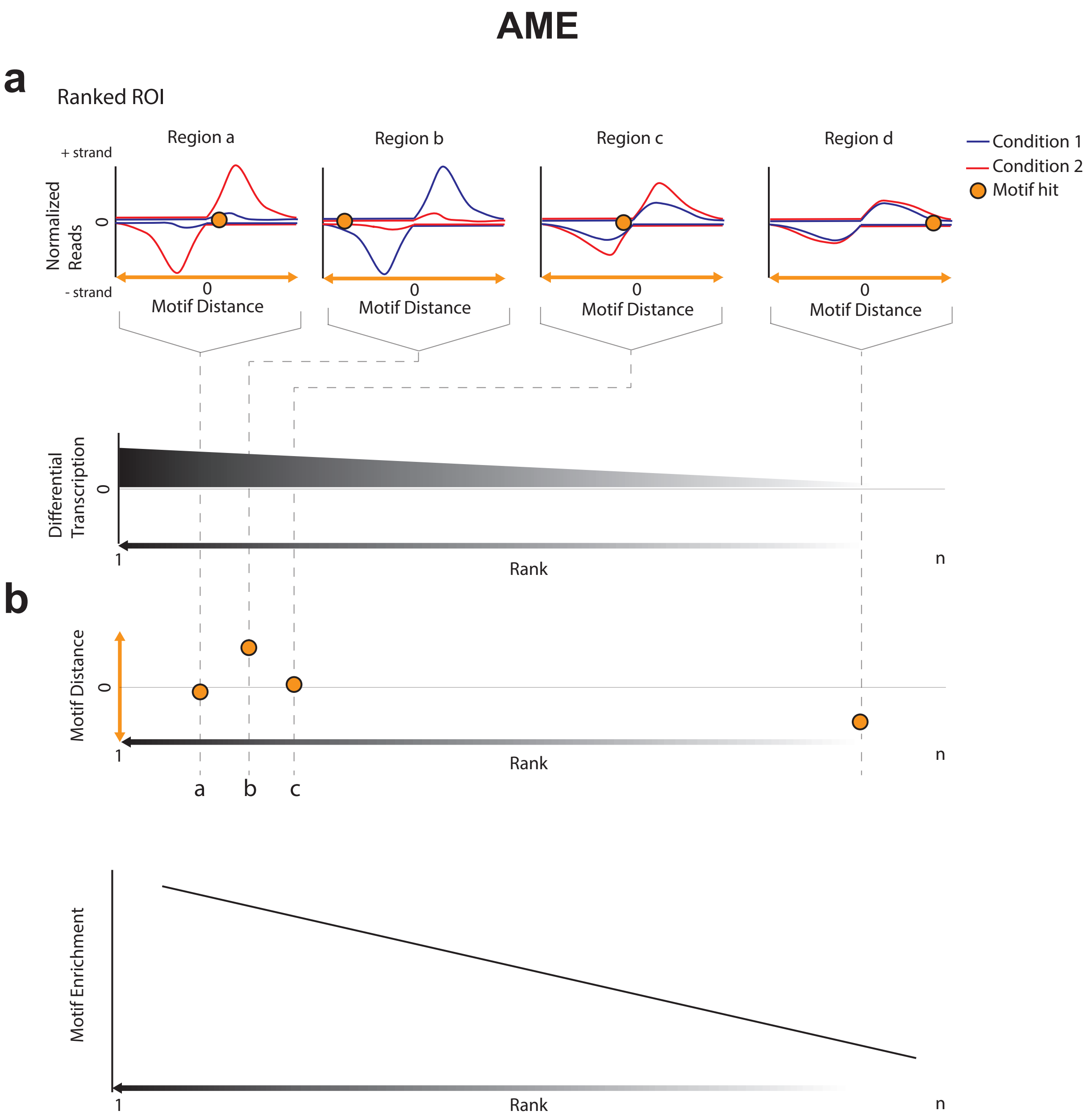


Supplemental Figure 13: **A cartoon diagram depicting the AME method.** Analysis of Motif Enrichment (AME) is part of the MEME suite and requires (a) a ranked list of ROIs as input. AME then performs (b) linear regression on the motifs as a function of rank.

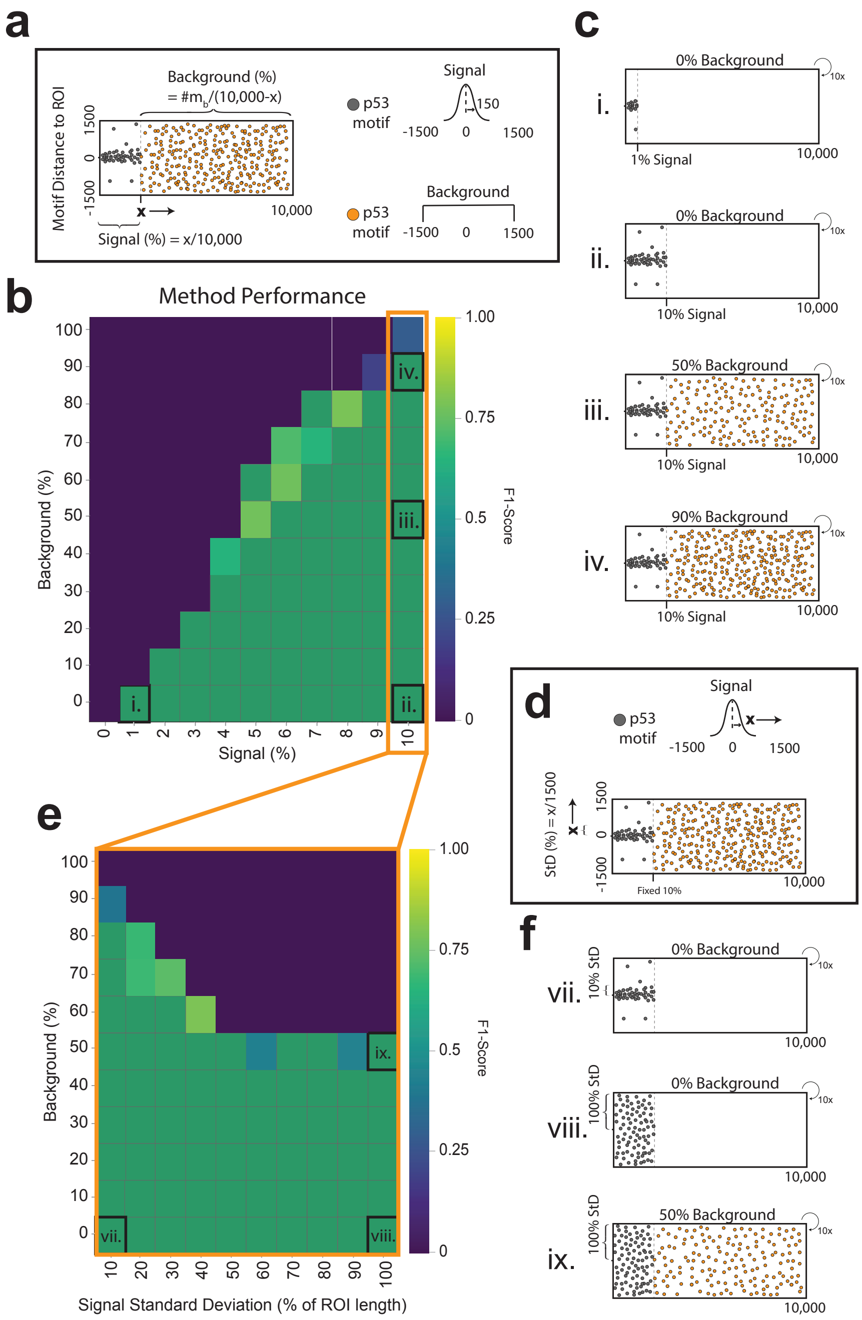


Supplemental Figure 14: **Diagram depicting the benchmark strategy utilized in Figure 4.** (a) A description of key concepts of motif embedding strategy for both signal (grey) and background (orange). (b) F1-Score (as heatmap) for benchmark varying fraction of ROI with signal (x-axis) and background (y-axis). Representative tests cases are labeled (i-iv) and their (c) respective embedding strategies are shown. (d) A description of additional criteria utilized for altering variability of signal embedding. (e) For 10% signal, we additionally alter the signal standard deviation (x-axis) vs background (y-axis). Representative cases (vii-ix) are labeled and their (f) respective embedding strategies are shown.

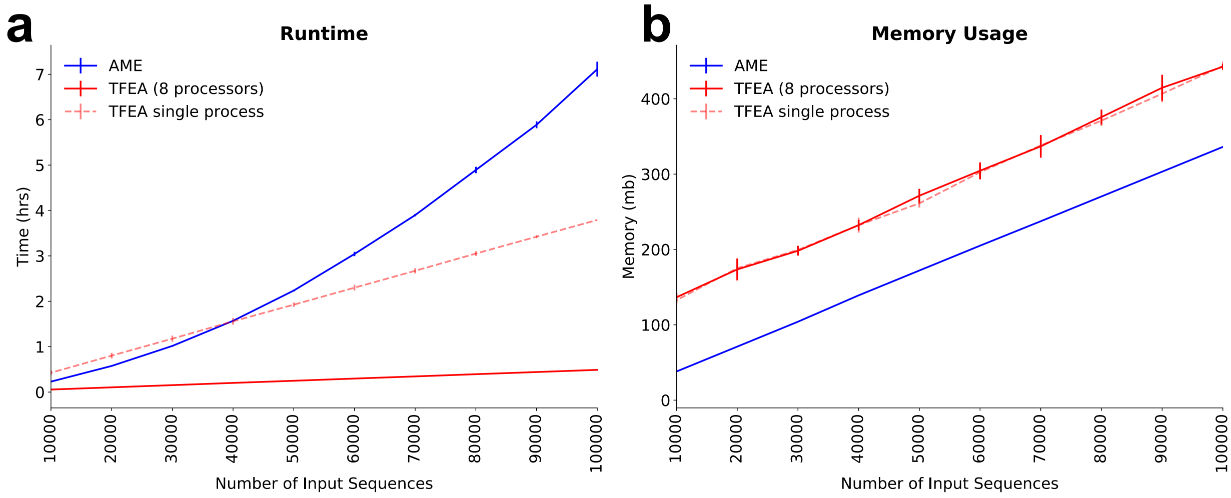


Supplemental Figure 15: **TFEA is fast and memory efficient.** (a) Runtime statistics for AME (solid blue; parallel processing not supported) and TFEA (8 processors: solid red; 1 processor: dashed red) with varying numbers of input ROI (bars = standard deviation of 10 runs). (b) Memory usage statistics comparing AME to TFEA.

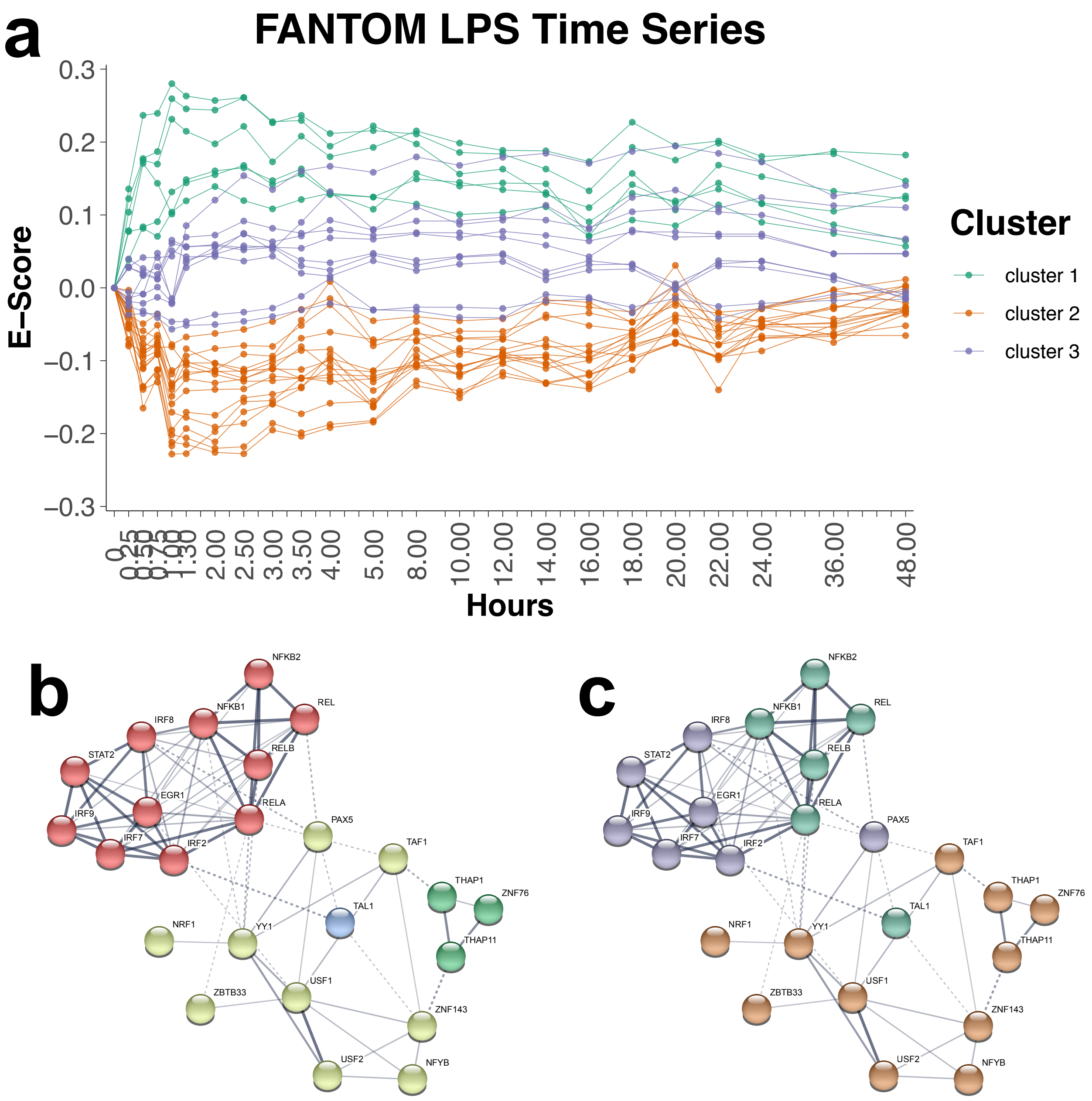


Supplemental Figure 16: **Clustering LPS induced TFs based on dynamics over time.** We applied k-means clustering to the subset of TFs that were significant (by TFEA) in at least 15 time points (~2/3 of all timepoints; n=32 TFs). (a) Time series traces of significant TFs colored by resulting cluster. The three main clusters correspond to the immediate increased responders (cluster 1, green), the immediate decreased responders (cluster 2, orange) and the later responding TFs (cluster 3, purple). (b) Alternatively the TFs can be analyzed using the String database using the Markov cluster algorithm. (c) Superposition of the coloring scheme in (a) onto the network cluster of (b).

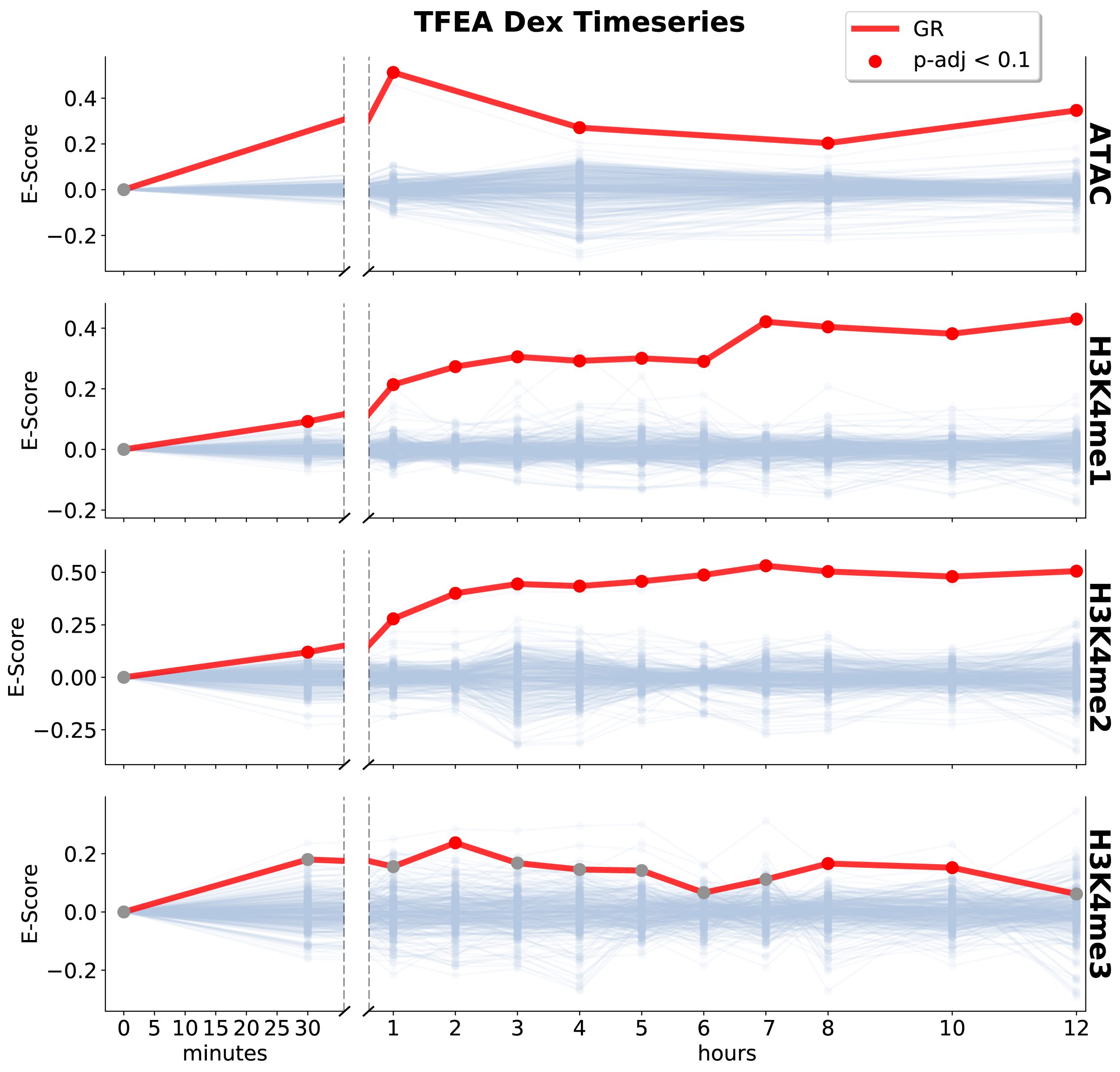


Supplemental Figure 17: **TFEA recovers the glucocorticoid receptor (GR) following treatment with Dexamethasone.** TFEA is able to recover GCR in many distinct data sets including ATAC, H3K4me1, and H3K4me2. Interestingly, TFEA only detects moderate enrichment of GR in H3K4me3, in agreement which previous results indicating that GR primarily binds to enhancers (which do not have the H3K4me3 mark).

[6] A. R. R. Forrest, H. Kawaji, M. Rehli, J. Kenneth Baillie, M. J. L. de Hoon, V. Haberle, T. Lassmann, I. V. Kulakovskiy, M. Lizio, M. Itoh, R. Andersson, C. J. Mungall, T. F. Meehan, S. Schmeier, N. Bertin, M. Jørgensen, E. Dimont, E. Arner, C. Schmidl, U. Schaefer, Y. A. Medvedeva, C. Plessy, M. Vitezic, J. Severin, C. A. Semple, Y. Ishizu, R. S. Young, M. Francescatto, I. Alam, D. Albanese, G. M. Altschuler, T. Arakawa, J. A. C. Archer, P. Arner, M. Babina, S. Rennie, P. J. Balwierz, A. G. Beckhouse, S. Pradhan-Bhatt, J. A. Blake, A. Blumenthal, B. Bodega, A. Bonetti, J. Briggs, F. Brombacher, A. Maxwell Burroughs, A. Califano, C. V. Cannistraci, D. Carbajo, Y. Chen, M. Chierici,

Y. Ciani, H. C. Clevers, E. Dalla, C. A. Davis, M. Detmar, A. D. Diehl, T. Dohi, F. Drabløs, A. S. B. Edge, M. Edinger, K. Ekwall, M. Endoh, H. Enomoto, M. Fagiolini, L. Fairbairn, H. Fang, M. C. Farach-Carson, G. J. Faulkner, A. V. Favorov, M. E. Fisher, M. C. Frith, R. Fujita, S. Fukuda, C. Furlanello, M. Furuno, J.-i. Furusawa, T. B. Geijtenbeek, A. P. Gibson, T. Gingeras, D. Goldowitz, J. Gough, S. Guhl, R. Guler, S. Gustincich, T. J. Ha, M. Hamaguchi, M. Hara, M. Harbers, J. Harshbarger, A. Hasegawa, Y. Hasegawa, T. Hashimoto, M. Herlyn, K. J. Hitchens, S. J. Ho Sui, O. M. Hofmann, I. Hoof, F. Hori, L. Huminiecki, K. Iida, T. Ikawa, B. R. Jankovic, H. Jia, A. Joshi, G. Jurman, B. Kaczkowski, C. Kai, K. Kaida, A. Kaiho, K. Kajiyama, M. Kanamori-Katayama, A. S. Kasianov, T. Kasukawa, S. Katayama, S. Kato, S. Kawaguchi, H. Kawamoto, Y. I. Kawamura, T. Kawashima, J. S. Kempfle, T. J. Kenna, J. Kere, L. M. Khachigian, T. Kitamura, S. Peter Klinken, A. J. Knox, M. Kojima, S. Kojima, N. Kondo, H. Koseki, S. Koyasu, S. Krampitz, A. Kubosaki, A. T. Kwon, J. F. J. Laros, W. Lee, A. Lennartsson, K. Li, B. Lilje, L. Lipovich, A. Mackay-sim, R.-i. Manabe, J. C. Mar, B. Marchand, A. Mathelier, N. Mejhert, A. Meynert, Y. Mizuno, D. A. de Lima Morais, H. Morikawa, M. Morimoto, K. Moro, E. Motakis, H. Motohashi, C. L. Mummery, M. Murata, S. Nagao-Sato, Y. Nakachi, F. Nakahara, T. Nakamura, Y. Nakamura, K. Nakazato, E. van Nimwegen, N. Ninomiya, H. Nishiyori, S. Noma, T. Nozaki, S. Ogishima, N. Ohkura, H. Ohmiya, H. Ohno, M. Ohshima, M. Okada-Hatakeyama, Y. Okazaki, V. Orlando, D. A. Ovchinnikov, A. Pain, R. Passier, M. Patrikakis, H. Persson, S. Piazza, J. G. D. Prendergast, O. J. L. Rackham, J. A. Ramilowski, M. Rashid, T. Ravasi, P. Rizzu, M. Roncador, S. Roy, M. B. Rye, E. Saijyo, A. Sajantila, A. Saka, S. Sakaguchi, M. Sakai, H. Sato, H. Satoh, S. Savvi, A. Saxena, C. Schneider, E. A. Schultes, G. G. Schulze-Tanzil, A. Schwegmann, T. Sengstag, G. Sheng, H. Shimoji, Y. Shimoni, J. W. Shin, C. Simon, D. Sugiyama, T. Sugiyama, M. Suzuki, N. Suzuki, R. K. Swoboda, P. A. C. 't Hoen, M. Tagami, N. Takahashi, J. Takai, H. Tanaka, H. Tatsukawa, Z. Tatum, M. Thompson, H. Toyoda, T. Toyoda, E. Valen, M. van de Wetering, L. M. van den Berg, R. Verardo, D. Vijayan, I. E. Vorontsov, W. W. Wasserman, S. Watanabe, C. A. Wells, L. N. Winteringham, E. Wolvetang, E. J. Wood, Y. Yamaguchi, M. Yamamoto, M. Yoneda, Y. Yonekura, S. Yoshida,

S. E. Zabierowski, P. G. Zhang, X. Zhao, S. Zucchelli, K. M. Summers, H. Suzuki, C. O. Daub, J. Kawai, P. Heutink, W. Hide, T. C. Freeman, B. Lenhard, V. B. Bajic, M. S. Taylor, V. J. Makeev, A. Sandelin, D. A. Hume, P. Carninci, Y. Hayashizaki, and The FANTOM Consortium and the RIKEN PMI and CLST (DGT). A promoter-level mammalian expression atlas. *Nature*, 507(7493):462–470, Mar. 2014.

[7] M. A. Gruca, M. A. Gohde, and R. D. Dowell. Annotation agnostic approaches to nascent transcription analysis: Fast read stitcher and transcription fit. *Methods in Molecular Biology*, to appear, 2019.

[8] I. C. McDowell, A. Barrera, A. M. D'Ippolito, C. M. Vockley, L. K. Hong, S. M. Leichter, L. C. Bartelt, W. H. Majoros, L. Song, A. Safi, D. D. Koçak, C. A. Gersbach, A. J. Hartemink, G. E. Crawford, B. E. Engelhardt, and T. E. Reddy. Glucocorticoid receptor recruits to enhancers and drives activation by motif-directed binding. *Genome Research*, Aug. 2018.

[9] S. K. Sasse, M. Gruca, M. A. Allen, V. Kadiyala, T. Song, F. Gally, A. Gupta, M. A. Pufall, R. D. Dowell, and A. N. Gerber. Nascent transcript analysis of glucocorticoid crosstalk with tnf defines primary and cooperative inflammatory repression. *Genome Research*, 2019.
